# Molecular height measurement by cell surface optical profilometry (CSOP)

**DOI:** 10.1101/2019.12.31.892075

**Authors:** Sungmin Son, Sho C. Takatori, Brian Belardi, Marija Podolski, Matthew H. Bakalar, Daniel A. Fletcher

## Abstract

The physical dimensions of proteins and glycans on cell surfaces can critically affect cell function, for example by preventing close contact between cells and limiting receptor accessibility. However, high-resolution measurements of molecular heights on native cell membranes have been difficult to obtain. Here we present a simple and rapid method that achieves nanometer height resolution by localizing fluorophores at the tip and base of cell surface molecules and determining their separation by radially averaging across many molecules. We use this method, which we call cell surface optical profilometry (CSOP), to quantify height of key multi-domain proteins on a model macrophage and cancer cell, as well as to capture average protein and glycan heights on native cell membranes. We show that average height of a protein is significantly smaller than its contour length due to thermally driven bending and rotation on the membrane and that height strongly depends on local surface and solution conditions. We find that average height increases with cell surface molecular crowding, while it decreases with solution crowding by solutes, both of which we confirm with molecular dynamics simulations. We also use experiments and simulations to determine the height of an epitope based on the location of an antibody, which allows CSOP to profile various proteins and glycans on a native cell surface using antibodies and lectins. This versatile method for profiling cell surfaces has the potential to advance understanding of the molecular landscape of cells and its role in cell function.

## Introduction

The surface of cells contains a dense and diverse population of molecules including proteins, glycosylated proteins, and glycolipids that extend above the plasma membrane. The average height of these proteins and glycans, which reflects their most common configurations, depends on structure as well as flexibility and local interactions. The size of individual folded domains can be obtained from crystal structures, which provide good estimates for the height of membrane proteins with single ectodomains, such as the 5 nm tall ‘marker of self’ CD47 (1). However, membrane proteins with ectodomains larger than a single IgG domain, which constitute approximately 51% of human surface proteome (2) (see Materials and Methods), have heights that are not simply the sum of their individual domains due to the domain organization, flexibility between domains, and thermally-driven fluctuations that reduce average extension from the surface. For example, the average height of the 7-domain CEACAM5 adhesion molecule will be smaller than its maximum fully-extended height of 29.7 nm obtained by summing domain sizes. This uncertainty in multi-domain protein height is compounded by contributions from unstructured domains and glycosylation that make estimates based on sequence and structure alone imprecise. For the protein tyrosine phosphatase CD45, which has four structured domains, for example, the average height of its isoforms will be variable depending on an N-terminal unstructured domain, and multiple glycosylation sites. Moreover, at densities of ∼20,000 molecules per square micron on the cell surface (3), which corresponds to an average spacing of ∼7 nm between molecules, the average heights of membrane proteins and glycans may be further altered by crowding.

Recent studies have revealed a critical role for cell surface molecular landscapes in cell function. In the immune system, size-dependent segregation of proteins has been shown to drive T-cell activation by antigen-presenting cells (4–7), broaden mast cell and basophil reactivity (8), and promote antibody-dependent phagocytosis by macrophages (9), where an antigen height difference of as little as 5 nm can significantly alter phagocytic efficiency. In cancer, increased glycocalyx height can drive integrin over-activation and promote tumor cell survival (10) and metastasis (11). In simulations of integrin binding, an increase in the glycocalyx height of 4 nm was predicted to reduce integrin bound fraction by an order of magnitude (12), while simulations of fluctuating membranes predicted that nanometer-scale height displacements of proteins can significantly alter ligand-receptor binding kinetics (13).

Despite the importance of the cell surface molecular landscapes in diverse cell functions, current techniques are not able to provide rapid and specific nanometer-scale measurements of cell surface molecular heights on native cell membranes. Transmission electron microscopy has been used extensively to image the glycocalyx of cells (14), though it is limited in its molecular specificity and requires lengthy sample preparations known to distort the glycocalyx (15). Several optical-based methods have been developed to quantify distances, such as scanning-angle interference microscopy (16) and fluorescence interference contrast microscopy (17, 18), but these methods require close contact of cells with a surface that can alter the extension of cell surface molecules. Fluorescence resonance energy transfer (FRET) can be used to quantify distances between two fluorophores on a single molecule (19), though FRET-based approaches lack the dynamic range needed to capture the full range of cell surface molecular heights, which can extend 300 nm above the membrane for some glycosylated proteins (14). Three-dimensional super-resolution imaging techniques such as iPALM (20), point spread function engineering (21), and multiplane imaging (22, 23) have been successfully used to image protein organization in integrin-(24) and cadherin-based cell adhesion (25), as well as to quantify the average thickness of the glycocalyx of cancer cells (26). However, these techniques often require a cell to be fixed to achieve high resolution, which can disrupt the native configurations of cell surface molecules, and typically do not exceed single-molecule localization accuracy of ∼10 nm resolution, unless highly specialized (27, 28). Since cell surface molecular height differences of just a few nanometers can significantly change cell function, a simple characterization method with resolution significantly better than 10 nm is needed.

Here we present a method for quantifying cell surface molecular height with ∼1 nm resolution that can be used to rapidly profile proteins and glycans on unfixed membrane surfaces, including native cell membranes. This method, which we call Cell Surface Optical Profilometry (CSOP), uses radial averaging of thousands of fluorescently-labeled molecules to achieve high localization accuracy and nanometer-scale height resolution (Fig. 1a). We validate CSOP by measuring the average end-to-end distance of tethered double-stranded DNA and by quantifying the thickness of bilayer membranes formed from different lipid compositions. We then use CSOP to measure the height and determine the flexibility of native and synthetic multi-domain proteins, and we investigate how crowding and solution conditions can alter height. Molecular dynamics simulations used to corroborate CSOP measurements reveal that certain dynamical/physical features of cell surface proteins may be understood in the classical framework of semiflexible polymers. We show that CSOP can also be used to localize the protein domain targeted by an antibody and to determine the average heights of individual molecular species as well as the entire cell surface proteome and glycome on native cell membranes. This simple, rapid, and versatile method for height measurement can provide new insight into the role played by cell surface molecular landscapes in diverse cellular processes.

**Figure 1:**
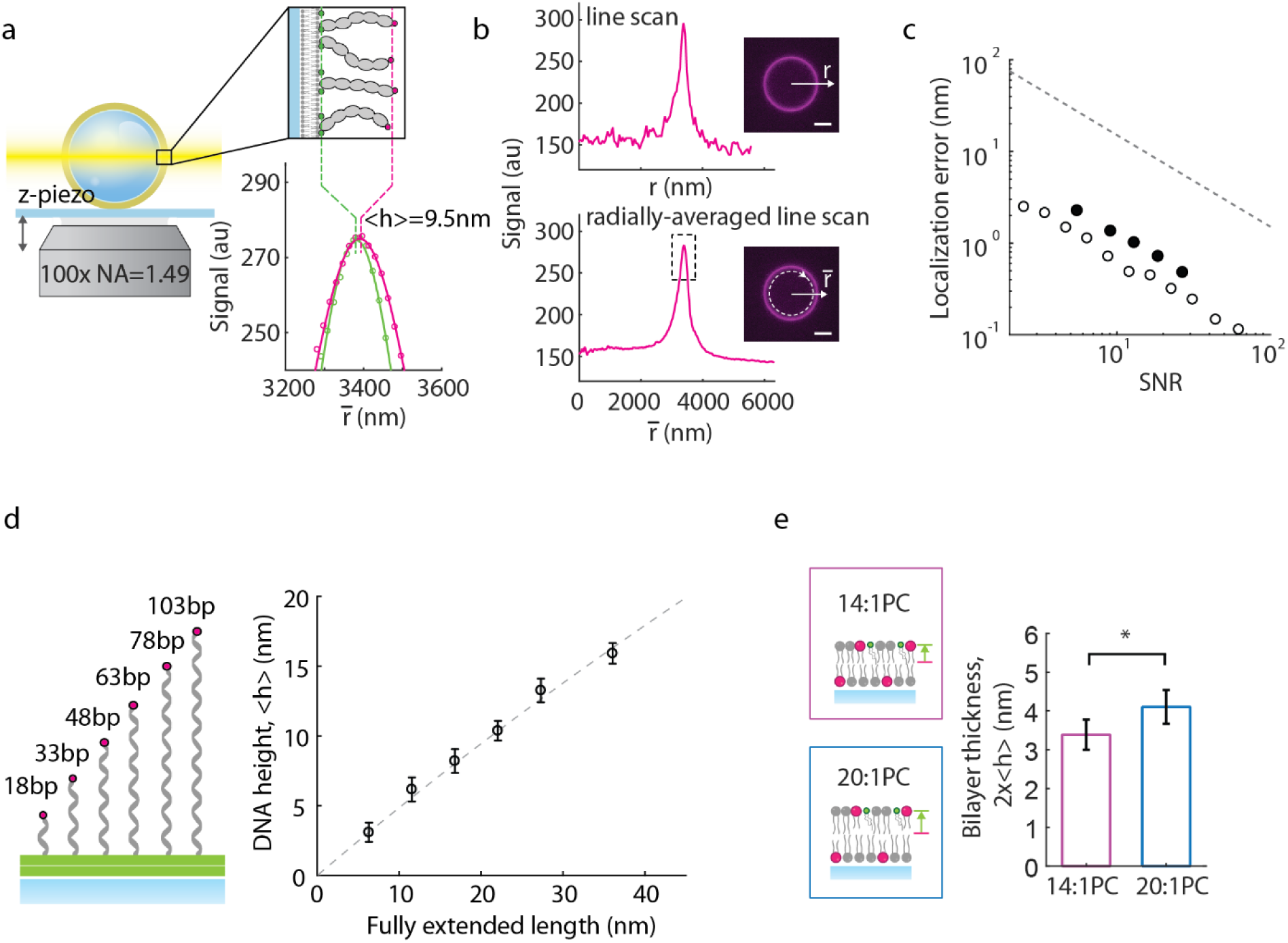
Cell Surface Optical Profilometry (CSOP) measures cell surface molecular heights using two-point localization: a) In an example CSOP measurement, a lipid-coated glass bead with multi-domain proteins bound to the membrane is imaged using confocal microscopy with a high-NA objective while a z-piezo stage scans the bead through a confocal plane. Inset shows a bilayer labeled with green fluorescent dyes and multi-domain proteins labeled with red fluorescent dyes at their tip. Protein height, <h>, is measured by localizing the centroids of the green and red fluorescent peaks averaged axisymmetrically. The fluorescence intensities of the protein or lipid channels (red and green circles) and their corresponding Gaussian fits (red and green lines) are shown below. b) A representative fluorescent profile of a protein on a lipid-coated bead along a single line *r* (top) or by radial averaging *r̅* of fluorescence signal (bottom), showing the improved SNR with radial averaging. The dashed grey box in the bottom linescan is zoomed in (a). Scale bar=2 µm. c) Comparison of CSOP’s resolution (circles) to other single-molecule localization methods (grey dotted line). The open circles were obtained from simulated data and the closed circles were obtained from experimental data. d) CSOP measurement of surface-tethered dsDNAs of varying length. Dotted line indicates the predicted WLC height when the persistence length is 50 nm, showing good agreement. N>40 for all measurements. The error bar indicates standard deviation. e) Quantification of lipid bilayer thickness with CSOP. Inset illustrates the location of a green (TopFluor-cholesterol) and red (Liss-Rhod b) label within a bilayer. The magenta and blue bars show the measurement of a bilayer containing either 14:1PC or 20:1PC lipids, respectively (N=99 or 103). Error bars indicate the 95% confidence intervals. p-value is 0.015 based on two-sample Student’s t-test.

## Results

### The principle of CSOP

Cell Surface Optical Profilometry draws inspiration from recent super-resolution microscopy methods that are based on precise localization of individual flourophores (29). However, rather than localizing in three dimensions, CSOP achieves nanometer-scale resolution of cell surface molecules in one dimension, height, by foregoing positional information in lateral dimensions. Height is determined using two fluorophores, one at the tip of the cell surface molecule to be measured and the other on the membrane in which the molecule is anchored. The fluorescently-labeled molecules are arranged in a spherical geometry, such as on a glass bead, giant vesicle, or swollen cell, so that multiple molecules oriented axisymmetrically can be averaged to obtain a measurement of fluorophore radius with high precision. Height is then quantified as the difference in radii of the two fluorophores.

This approach overcomes the typical ∼10 nm resolution limit of single molecule localization by increasing the number of fluorophores used in the radius localization by orders of magnitude (Fig. 1b). Tens to hundreds of thousands of fluorophores indicating the same position in the radial coordinate enhances the radial intensity profile above background noise, enabling high-resolution determination of average separation between fluorophores. Typical STORM imaging collects 3,000 photons from a single fluorophore per exposure, which corresponds to a ratio of peak signal to photon shot noise of 20. In practice, this SNR = 20 enables localization of the fluorophore with 8 nm precision, after careful removal of drift (30). For a measurement of fluorescently-labeled proteins bound to the surface of a bead (Fig. 1a & 1b), CSOP achieves 0.65 nm precision for the same SNR by radially averaging, in this case, ∼550 labeled proteins/µm^2^ from which we collect ∼100 photons/fluorophore (Fig. 1b, Table S1). If we increase the density of labeled proteins but continue to collect the same number of photons/fluorophore, this localization precision is increased to 0.5 nm for 1000 labeled proteins/µm^2^, though this is below the predicted precision of 0.3 nm that should be theoretically possible for a shot noise limited measurement (Fig. 1c, Table S1). For typical densities of labeled molecules on cell surfaces, a height resolution of ∼1 nm is readily obtained.

Notably, this approach to cell surface molecular height measurement enables simple multiplexing of height measurements by labeling different cell surface molecules on the same membrane with different fluorophores, making it possible for CSOP to simultaneously profile multiple proteins and glycans using standard multi-channel fluorescence microscopy.

### Validation of molecular height measurements with CSOP

We set up CSOP on a standard spinning disk confocal microscope with a sCMOS camera and a high-NA objective (Yokogawa CSU-X1, Andor Zyla 4.2, 100×/1.49 Apo). To demonstrate height measurements with CSOP, we used lipid-coated glass beads with purified molecules attached to the outer leaflet of the membrane, with Alexa 488 localizing the membrane distal end of the molecule and Alexa 555 localizing the membrane surface. We corrected for chromatic aberrations, which can introduce errors into CSOP measurements (Fig. S1) (31), by implementing a calibration procedure before imaging on a new microscope (Fig. S2, Materials and Methods). We also addressed polarization effects associated with membrane inserting labels by using an alternate membrane-bound label (Fig. S3, Materials and Methods) (32).

As a validation of CSOP’s height measurement capabilities, we measured the end-to-end distance of double-stranded DNA tethered to a surface. We quantified the average separation between a bilayer membrane and the fluorescently-labeled end of double-stranded DNA ranging from 18 to 103 basepairs (Table S2). We found that the CSOP height measurements, <h>, agree well with predictions based on previously measured DNA end-to-end lengths and a persistence length of 50 nm (33) (Fig. 1d, Materials and Methods). These measurements show that the average height of short dsDNA tethered to a surface and able to freely rotate is approximately half of its fully extended length, and they also confirm CSOP’s ability to accurately capture a broad range of molecular heights.

As a test of CSOP’s resolution, we measured the thickness of lipid bilayers formed from either 14- or 20-carbon lipids. We compared the average position of one fluorophore labeling both leaflets of a supported lipid bilayer on a glass bead to the average position of a second fluorophore labeling only the outer leaflet (Fig. 1e and Materials and Methods). The difference in radius of the two different labeling schemes reports the thickness of one leaflet. Using this approach on approximately 100 beads composed of either 14- or 20-carbon lipids, we were able to detect a <h>= 0.36 +/- 0.1 nm (S.E.M.) for the thickness of lipid leaflets due to acyl chain length. This gives a difference in bilayer thickness (2×<h>) that is consistent with previous measurements by imaging ellipsometry (34) and confirms that sub-nanometer resolution is possible with CSOP.

### Measurement of multi-domain protein height and flexibility with CSOP

To demonstrate the ability of CSOP to measure nanometer-scale differences in the height of multi-domain proteins, we constructed a series of FN3 domain repeat proteins (9). The FN3 proteins ranged from a single domain (1L) to eleven domains (11L), with molecular weights ranging from 14 kDa to 154 kDa, respectively. We attached the purified multi-FN3-domain proteins via a His-tag to supported lipid bilayers containing Ni-chelating lipids on a glass bead. We labeled each protein with an N-terminal ybbR-tag, and we labeled the bilayer with an AlexaFluor dye-conjugated His-tag peptide (Fig. 2a, Materials and Methods). Using CSOP, we measured the heights of each of the multi-FN3-domain proteins and compared it with the estimated fully-extended height, which was obtained by multiplying the FN3 domain size based on its crystal structure by the number of domains. As expected for flexible, multi-domain proteins, the measured height was consistently shorter than the proteins’ maximum fully-extended height (Fig. 2a).

**Figure 2:**
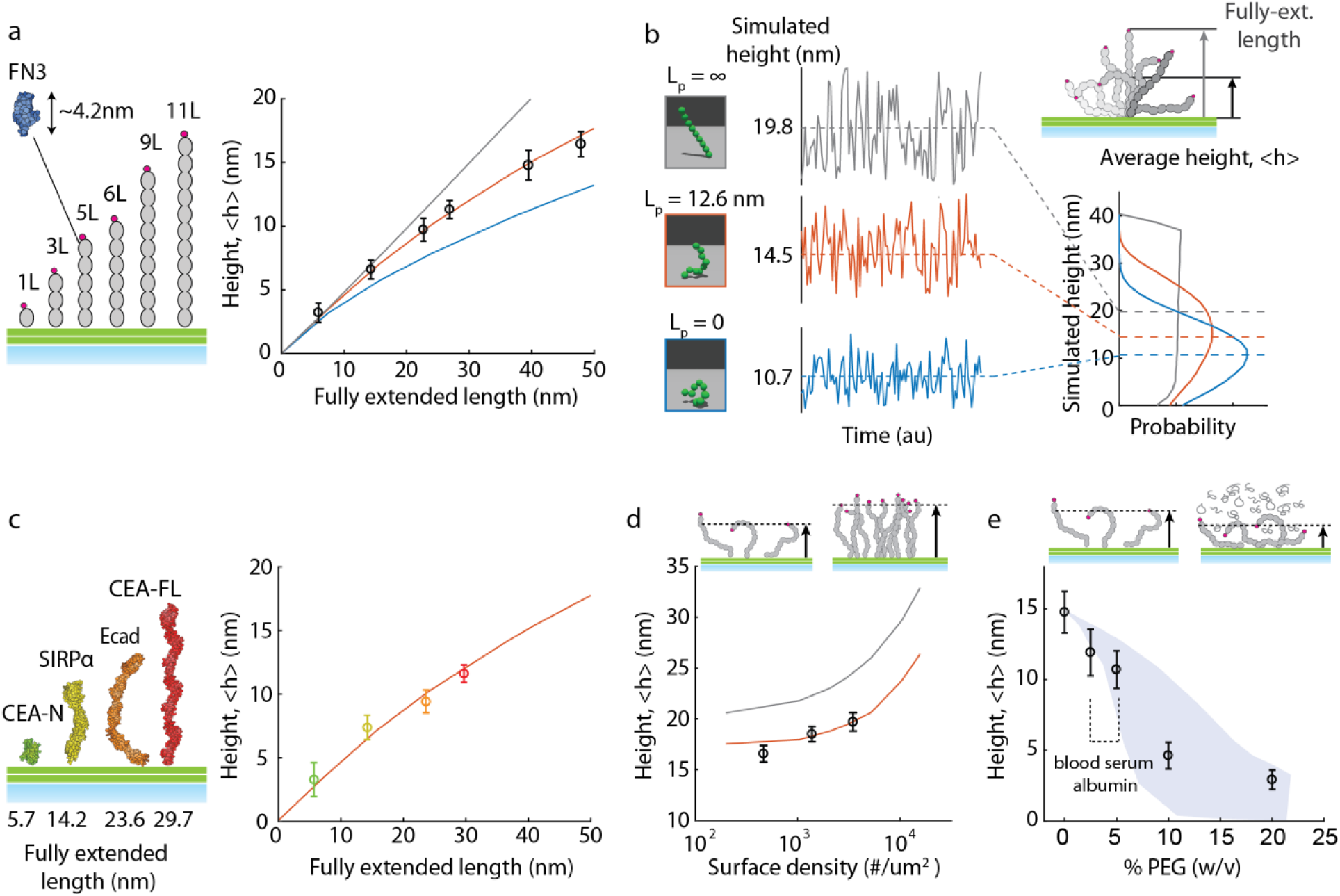
CSOP measurements of multi-domain proteins’ height and flexibility: a) CSOP height measurements of a series of multi-FN3-domain proteins. The lines indicate molecular dynamics (MD) simulation of a rigid-rod (grey), a semi-flexible polymer with the persistent length 12.6 nm (red), and a freely-jointed chain (blue). The size of a single FN3 domain is assumed to be 4.2nm based on its structure (PDB 3TEU). N>25 for all proteins. b) MD simulations of a 9L FN3-domain protein height with different persistence lengths vs. time. The dotted lines indicate the mean height. The graphics show an instantaneous protein configuration. Top-right: illustration of a protein on a membrane surface while bending and rotating. Bottom-right: Protein height probability distributions of the 9L FN3-domain based on MD simulations. c) Height measurements of native cell surface proteins human CEACAM5 N-terminal domain (CEA-N), mouse SIRPα (PDB 2wng), mouse E-cadherin (PDB 3q2v), and human CEACAM5 full-length protein (CEA-FL; PDB 1e07). Fully extended lengths are based on crystal structures. The red line indicates an MD simulation of a semi-flexible polymer with persistence length 12.6 nm, same as in a). N>25 for all measurements. d) Average height measurements of 11L FN3-domain protein at three different surface densities (10 nM, 40 nM, and 180 nM in solution). N=92, 50, and 79, respectively. The curves show MD simulation results for a rigid-rod (grey) and a semi-flexible polymer with the persistence length 12.6 nm (red). e) Average height of 9L FN3-domain protein is measured with varying concentrations of PEG8K in solution. The dotted line indicates physiological concentration of blood serum albumin. The shaded area shows MD simulation results when the radius of gyration of PEG8k was varied from 2.6 nm to 3.5 nm. N >15 for all measurements. The error bars in all figures indicate standard deviation.

We next sought to use our measurement of average height to determine the flexibility of the multi-FN3-domain proteins using course-grained molecular dynamics (MD) simulations. We simulated proteins tethered to membranes using a worm-like chain model with a defined persistence length (L_p_), which captures the resistance to bending, with L_p_=0 representing a freely-jointed chain (blue) and L_p_=infinity representing a rigid rod (grey) (Figs. 2b, Materials and Methods, Figs. S4-6, Movie S1). Comparing the simulations to the CSOP data, we found that the height measurements for all multi-FN3-domain proteins were consistent with simulations of a semi-flexible polymer, L_p_∼12.6 nm (red), indicating that the orientation of the synthetic multi-FN3-domain proteins is highly correlated (rod-like) over approximately three domains.

We then used CSOP to measure the height of a series of native multi-domain cell surface proteins whose average height and flexibility are not known (Fig. 2c). Specifically, we measured mouse signal-regulatory protein alpha (SIRPα) (three Ig domains), mouse E-cadherin (five cadherin domains), and human carcinoembryonic antigen related cell adhesion molecule 5 (CEACAM5) full-length protein (seven Ig-like domains), as well as the N-terminal domain of CEACAM5 alone (one Ig-like domain). For the extracellular domain of each protein, we added an N-terminal ybbR-tag for fluorescent labeling and a C-terminal His-tag for membrane binding. Similar to our measurement of multi-FN3-domain proteins, all four proteins behaved as worm-like chains with average heights shorter than their fully-extended height. Interestingly, all measured proteins were well described by L_p_∼12.6 nm, suggesting similar flexibility of these proteins and the multi-FN3-domain proteins. These data show that CSOP measurements combined with simulations can provide critical information on the height and flexibility of cell surface proteins that cannot be obtained from crystal structures.

### Effect of surface and solution crowding on multi-domain protein height

We next used CSOP to measure the effect of surface density on protein height. Molecular crowding of proteins on a membrane surface can create significant lateral pressure (35) that can alter the height of multi-domain proteins by constraining their fluctuations, as in the case of densely crowded MUC1 glycoproteins on the surface of epithelial cells (10). To quantify the effect of protein crowding on height with CSOP, we measured our longest multi-FN3-domain protein (11L) at three different surface densities. We found that the average height of the protein increased significantly, by 1.9 nm (11%) and 3.1 nm (19%), as surface concentrations increased by a factor of 3x and 7.5x, respectively, from an initial density of ∼460/µm^2^ (Fig. 2d). This change is consistent with our MD simulation of proteins on a cell surface for those densities, which further predicts that protein height can increase by up to 50% at surface densities equivalent to that of native cell membranes (>10,000/µm^2^) (3) (Materials and Methods).

Cell surface proteins not only experience lateral crowding from neighboring membrane-bound proteins but also macromolecular solution crowding from the surrounding medium. We used CSOP to detect the effect of solution crowding on height by measuring a long multi-FN3-domain protein (9L) under varying amounts of 8kDa polyethylene glycol (PEG) in solution. Interestingly, we found that protein height decreased by 20-30% for solution crowding concentrations of 3.5-5% w/v, which are physiological levels of solution crowding concentration (36), with more dramatic height decreases for solution crowding concentrations exceeding 5% w/v (Fig. 2e). Again, our measurements agreed well with results from MD simulations of solution crowding, which also predict that the absolute height decrease by bulk osmotic compression is larger for taller proteins than for shorter ones (Materials and Methods, Movie S2). Our data suggest that physiological concentrations of macromolecules in solution will not only affect the height of the thick glycocalyx (100-500nm) of epithelial (37) or endothelial (38) cells but also the height of cell surface proteins on nearly all cells.

### Measurement of protein height using antibodies

While CSOP requires a fluorescent marker at the tip of a cell surface molecule to measure its height, that marker need not be an engineered fluorescent tag. Indeed, for CSOP to be broadly useful for measuring cell surface proteins on primary cells, such as patient samples, an approach that does not require modification of native proteins is needed. Here we demonstrate the use of fluorescent antibodies targeting the terminal domain of surface proteins to enable protein height measurements on native cell membranes.

To validate antibody labeling, we first measured the heights of CEACAM5, SIRPα, and multi-FN3-domain proteins with antibodies targeting their terminal domains and compared those to measurements of the same proteins with engineered fluorescent tags on their terminal domains (Fig. 3a). We observed that antibodies add an offset of 1.5 +/- 1.4 nm (S.D.) to CEACAM5 and the offset increases for shorter proteins, presumably because the antibody is itself a relatively large molecule (∼7 nm for the Fab). We used MD simulations of antibody-protein complex motion to quantify how much size increase would be expected and found that steric effects result in an increase in apparent height of ∼1.6 nm (Materials and Methods, Fig. S7, Movie S3). This offset increases as protein size decreases due to surface effects, which is consistent with our experimental measurements with CSOP (Fig. 3a). The same trend in offset was observed regardless of a target protein’s flexibility or surface density in the simulations, suggesting that protein height can be accurately measured with a fluorescently-labeled antibody by subtracting the height-dependent offset added by the antibody.

**Figure 3:**
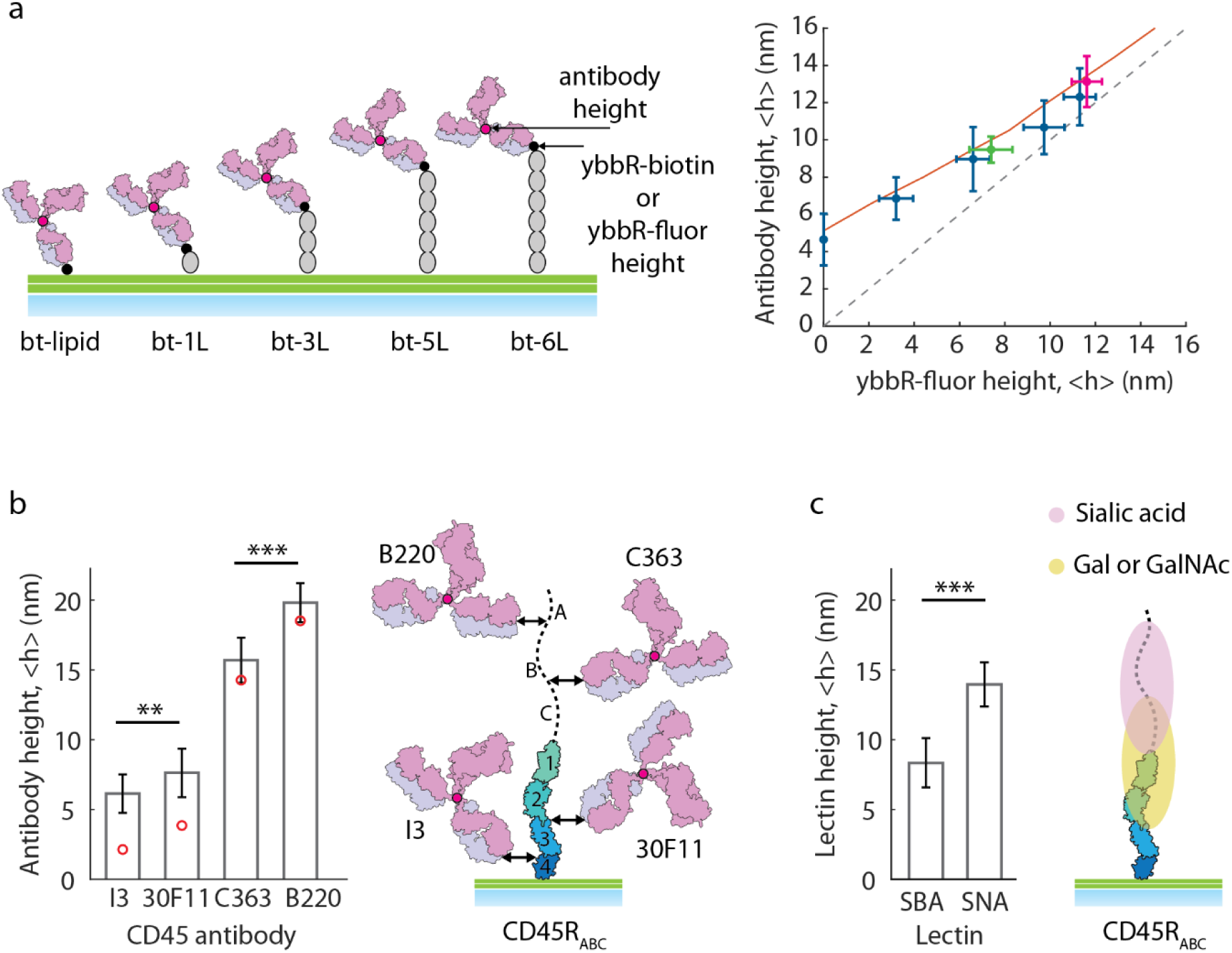
CSOP measurements of protein and glycan heights using fluorescent antibodies and lectins: a) Biotinylated (ybbr-biotin) multi-FN3-domain proteins (bt-1L to bt-6L) were labeled with fluorescently-labeled anti-biotin antibodies, as was a biotinylated lipid (bt-lipid). Black dots indicate biotin or engineered fluorescent tag and red dots indicate the antibody’s center-of-mass. Height measurements based on localizing the fluorescent antibody of the multi-FN3-domain proteins (y-axis, blue) are plotted against the protein height based on an engineered fluorescent tag (ybbR-fluor, x-axis), showing a size-dependent offset. Measurement of SIRPα (green) and CEACAM5 (magenta) are consistent with the multi-FN3-domain protein measurements. The red line indicates result from MD simulations, and the dotted line shows a case whereby the antibody location is identical to the protein height. N>25 for all measurements. b) Height measurement of antibodies bound to CD45R_ABC_. Red circles indicate epitope height after subtracting the offset added by the antibody. N>20 for all measurements. p-values based on two-sample Student’s t-test are 0.002 and 3.5×10^-11^. The measured heights of anti-CD45 antibodies are mapped onto a model of CD45R_ABC_, showing the predicted locations of the antibody epitopes (black arrows). c) Height measurement of lectins bound to CD45R_ABC_. N>20 for all measurements. All error bars indicate the standard deviation. p-value based on two-sample Student’s t-test is 4.2×10^-14^. The measured heights of glycans are mapped onto a model of CD45R_ABC_, showing a difference in the height of sialic acid and Gal/GalNAc residues.

### Measurement of antibody epitope location on a multi-domain protein

Identifying antibody epitopes usually requires co-crystallography, mutagenesis and binding measurements, or cross-linking-coupled mass spectrometry (39), all of which are time-consuming and costly methods. We wondered whether CSOP could be used to determine the approximate location of antibody epitopes on multi-domain proteins by measuring the antibody height and comparing it to protein height.

As a demonstration of epitope mapping with CSOP, we examined antibodies targeting CD45 – a transmembrane tyrosine phosphatase whose exclusion from membrane interfaces plays a critical role in immune cell activation (40, 41) (Fig. 3b). The CD45 extracellular domain is comprised of an N-terminal mucin-like region followed by a cysteine-rich domain (d1) and three fibronectin type 3 (FN3) (d2–d4). The longest human CD45 isoform, CD45R_ABC_, has a mucin-like region of 202 residues and is predicted to be approximately 40 nm when fully extended, with domains d1-d4 accounting for 15 nm of its height (41, 42).

We used four commercially available anti-CD45 antibodies that have different isoform specificities and are therefore thought to bind at different locations in CD45, though the specific epitopes and domain locations are unknown. We selected the pan-CD45 antibodies I3 and 30F11, the anti-CD45R_B_ antibody C363, and the anti-CD45R_ABC_ antibody B220. Using CSOP, we measured the height of the antibodies on CD45R_ABC_ bound to lipid-coated beads, and we were able to clearly separate the binding locations: I3 bound to the d4 domain (6.1 +/- 1.4 nm (S.D.)), 30F11 bound to the d2-d3 domain (7.6 +/- 1.7 nm (S.D.)), C363 bound to the proximal part of the mucin-like domain (15.7 +/- 1.6 nm (S.D.)), and B220 bound to the distal part of the mucin-like domain (19.8 +/- 1.4 nm (S.D.)) (Fig. 3b). Based on the measurement of the B220 antibody, the CD45R_ABC_ extends an average of 18.2 nm from the membrane surface, less than half the fully-extended height estimated from its structure (41, 42). These results suggest that antibody-binding sites separated by ∼1 nm within the same protein are resolvable using CSOP.

### Measurement of glycan height on a multi-domain protein

While we have focused on protein height measurements, the majority of cell surface proteins are glycosylated (43), the extent of which will alter their cell surface height. Importantly, glycosylation is not typically accounted for in crystal structures and is difficult to predict for a given protein, making direct measurement of glycans the most reliable way to determine their height. We tested CSOP’s ability to localize cell surface glycans using lectins that bind to specific glycans.

Using the purified extracellular domain of CD45R_ABC_, we measured the height of fluorescent plant lectins Soybean Agglutinin (SBA), which binds GalNAc and galactose, and Sambucus Nigra Lectin (SNA), which targets α-2,6 sialic acids, on CD45R_ABC_ (Fig. 3c). We found that the average height of SNA was 13.8 +/- 1.6 nm (S.D.), while the average height of SBA was 8.3 +/- 1.7 nm (S.D.). SNA was localized close to the height of the anti-CD45R_B_ antibody epitope, suggesting that CD45R_ABC_ is predominantly sialylated in the mucin-like domain. Our measurements demonstrate that CSOP can captures differences in the average position of specific glycans on the same molecule.

### Measurement of glycoprotein height on native cell membranes

We next applied CSOP to make high-resolution molecular height measurements on native cell membranes. While the use of antibodies to target endogenous cell surface proteins is compatible with CSOP, as demonstrated above, the arrangement of cell membranes in a spherical geometry is necessary to make use of radial averaging. Conveniently, several techniques have already been devised for preparing cells and cell membranes that are compatible with CSOP measurements, including giant plasma membrane vesicles (GPMVs) (44) and swollen cells (45). Despite some drawbacks arising from their preparation (46), these methods preserve the native composition of cell surface proteins, including their post-translational modifications, allowing CSOP measurement of protein heights in situ.

To test if CSOP can accurately measure molecular heights on giant plasma membrane vesicles (GPMV) (44), we compared the height of a protein we had previously measured on the supported bilayer of a lipid-coated glass bead, CEACAM5, to CSOP measurement of GPMVs from cells overexpressing CEACAM5 (Fig. 4a, Materials and Methods). GPMVs maintain the native lipid and protein diversity, allowing biochemical and biophysical studies of cell membrane and proteins separate from the cytoskeleton. We labeled CEACAM5 using a V_HH_ nanobody that recognizes a small peptide tag at its N-terminus (47). Based on CSOP height measurements, we found that CEACAM5’s height in GPMVs was consistent with its height measured in lipid-coated beads (Fig. 4b). We repeated GPMV measurements for the CEACAM5 N-terminal domain alone, as well as SIRPα and E-cadherin, all using the V_HH_ nanobody, and we found that average height measurements from the GPMVs were remarkably consistent with those from lipid-coated beads, except for SIRPα, which becomes 40% higher when measured in a GPMV (Fig. 4c). We wondered if the way in which SIRPα was attached to the membrane – with a transmembrane domain in native cell membranes and with a His-tag on the extracellular domain in vitro – could affect the measured height. To test the effect of membrane attachment, we swapped the wild-type transmembrane domain of SIPRα for a GPI-anchor. Interestingly, the height of SIPRα-GPI in GPMV exhibited only 27% increase from its height measured in lipid-coated beads, compared to 40% increase with the wild-type transmembrane domain (Fig. 4c). This suggests that the transmembrane anchor can significantly change protein height, in addition to other factors, showcasing the sensitivity of CSOP measurements for structural details of cell surface molecules.

**Figure 4:**
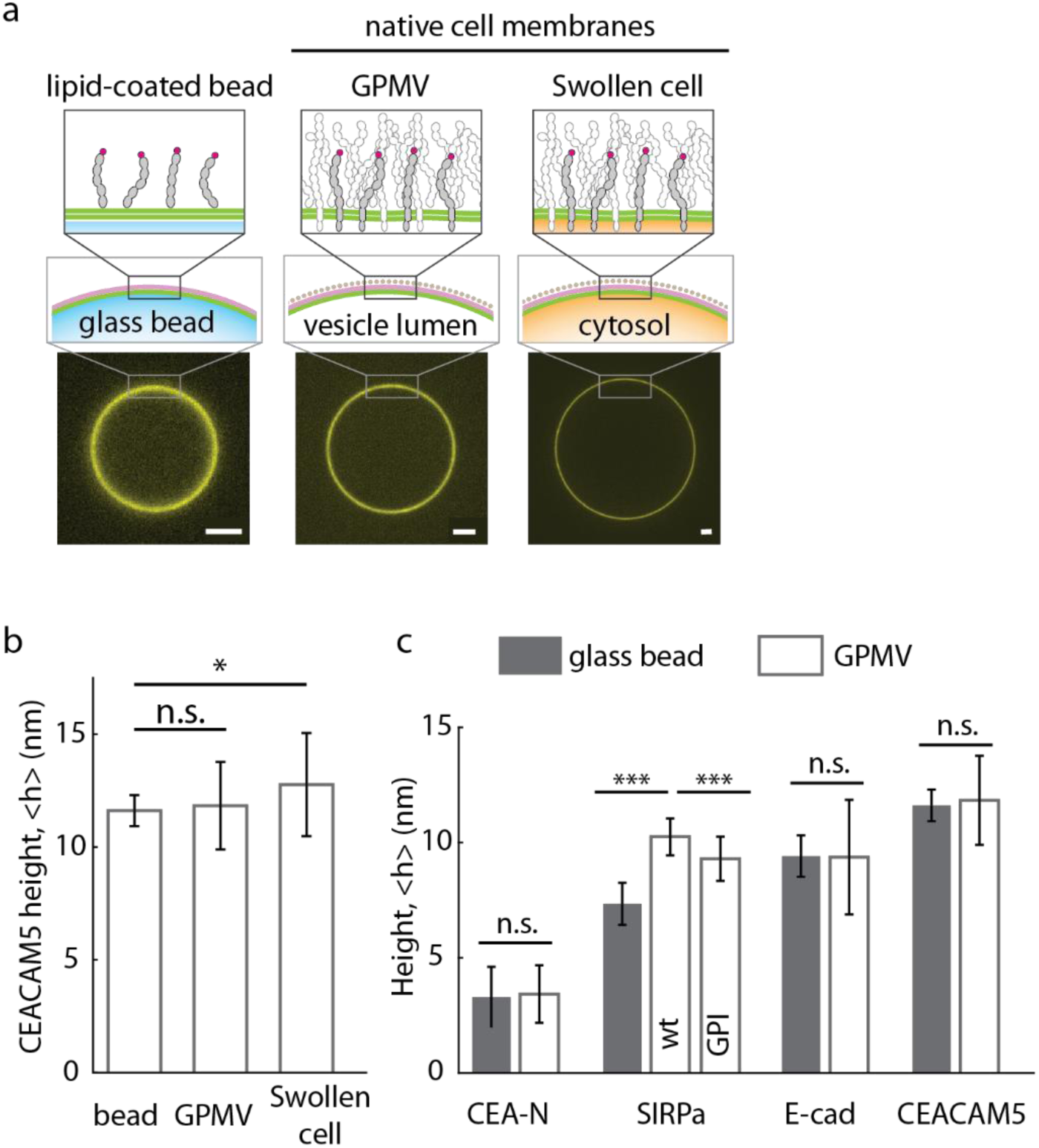
CSOP measurements of cell surface protein heights in native membranes: a) Illustration of protein and membrane preparations that can be measured with CSOP. Images show fluorescent CEACAM5 in the different preparations. Scale bar=2 µm. b) CSOP measurement of CEACAM5 in three different preparations. N=30, 40, and 26, respectively. p-values based on two-sample Student’s t-test are 0.56 and 0.011, respectively. c) Measurement of cell surface protein heights on lipid-coated beads (the closed columns) versus GPMVs (the open columns). p-values based on two-sample Student’s t-test are 0.63, 4.7×10^-19^, 7.0×10^-4^, 0.92, and 0.56, respectively. N>15 for all measurements.

Since some cell types are not amenable to GPMV formation, we adapted an existing protocol for osmotically swelling cells into spheres for use with CSOP (45) (Fig. 4a). To swell cells, we first disrupt the actin and microtubule cytoskeleton by using small molecule drugs, Latrunculin A and Y-27632 (Materials and Methods). Next, we hydrostatically swell cells by reducing the extracellular buffer osmolarity. More than 50% of HEK cells overexpressing CEACAM5 turned completely spherical at ∼80 mOsm and remained swollen for at least 1hr without any regulatory volume decrease (48). After cell swelling, we labeled the membrane using a CellMask dye and labeled CEACAM5 using the V_HH_ nanobody as above. Fluorescently-labeled CEACAMs were uniformly distributed on the membrane, making it indistinguishable from a GPMV. We then measured protein height and found that CEACAM5’s height on a swollen cell is ∼10% higher than its height measured on a lipid-coated bead (Fig. 4b).This is consistent with our previous observation that multi-FN3-domain proteins are more extended at high surface densities like those expected on a native cell membrane (Figure 2d).

### Measurement of proteome and glycome heights on native cell membranes

Height measurements based on antibodies and engineered tags can provide information on specific cell surface molecules, but it is often useful to know how tall the surrounding molecules are in comparison. This is particularly relevant when considering, for example, whether an antibody-bound antigen on a tumor cell is accessible to an immune receptor.

To explore this, we measured average cell surface proteome and glycome heights on a HER2-positive breast cancer cell, SKBR3, and on a macrophage progenitor cell, Hoxb8 (49). To measure average proteome height, we labeled the N-terminus of cell surface proteins using an amine-reactive biotin (NHS-Biotin) at low pH (pH 6.5)(50). We then swelled the cells, as described above, and measured the height of the cell surface biotin label using an anti-biotin antibody. The average cell surface proteome height of SKBR3 cells was measured to be 13.5 +/- 2.2 nm (S.D.) and that for the Hoxb8 cell was found to be 15 +/- 2.9 nm (S.D.). To determine the average cell surface glycome height, we used CSOP to measure the heights of GlcNAc and sialic acid with the lectins Wheat Germ Agglutinin (WGA) and Maackia Amurensis Lectin II (MAL-II), respectively. The average heights of α-2,3 sialic acid as measured by MAL-II were 16.4 +/- 3.7 nm (S.D.)(SKBR) and 16.7 +/- 2.6 nm (S.D.)(Hoxb8) (Fig. 5a and b). We also found that GlcNAc heights as measured by WGA were 7.8 +/- 1.9 nm (S.D.)(SKBR) and 5.9 +/- 1.8 nm (S.D.)(Hoxb8).

**Figure 5:**
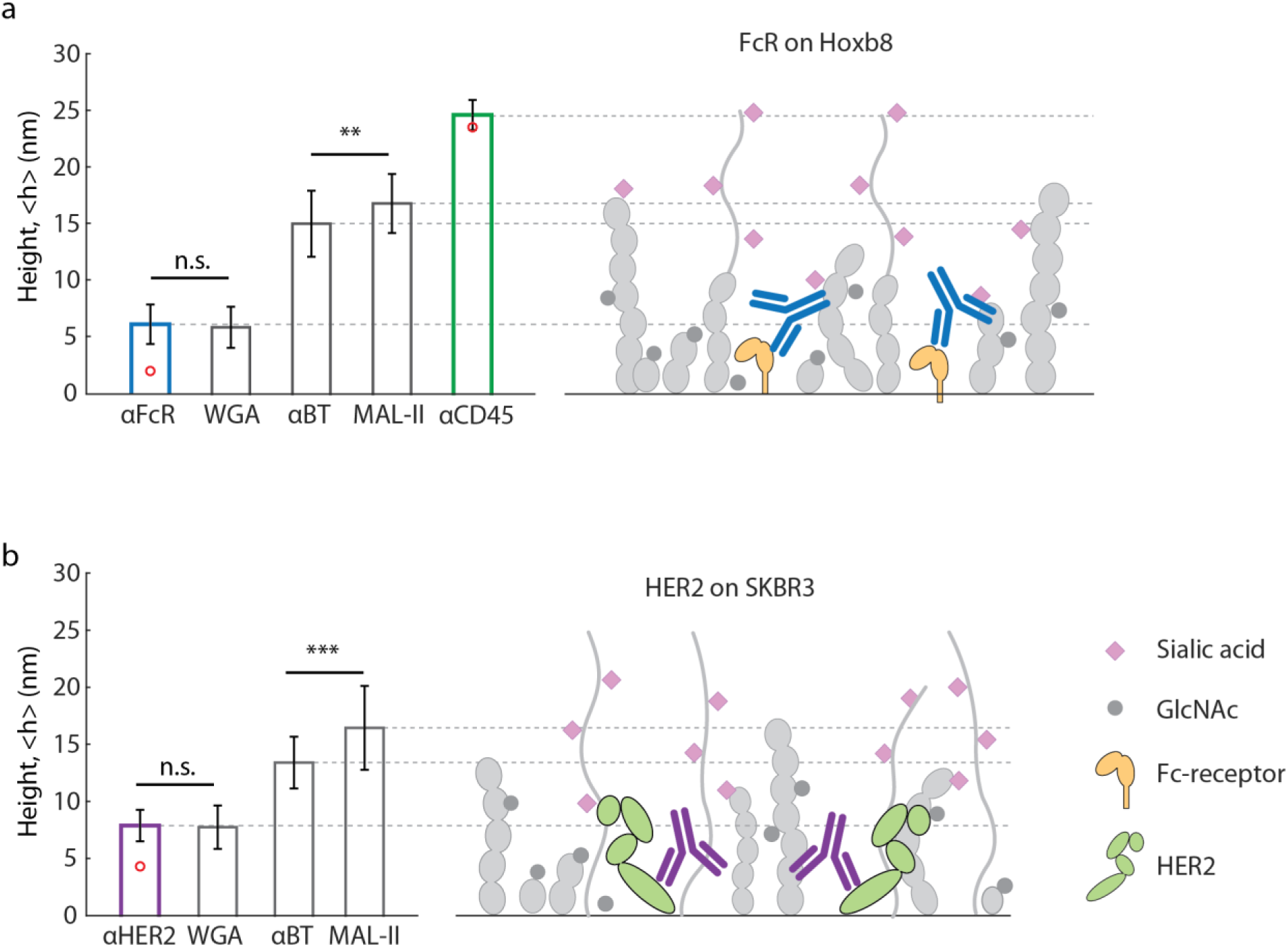
CSOP measurements of average cell surface proteome and glycome heights on tumor and immune cells: a) The height of Fc-receptor, CD45, average proteome, and average glycome were measured on Hoxb8 cells by binding antibodies or lectins. The results indicate that Fc-receptors are much shorter than the average proteome and some glycan structures. Anti-biotin, Anti-CD45, and lectins are not drawn. N>30 for all measurements. Error bars indicated standard deviation. p-values based on two-sample Student’s t-test are 0.66 and 5.0×10^-5^. b) The height of HER2, average proteome, and average glycome were measured on SKBR3 tumor cells by binding antibodies or lectins. The results indicate that Her2 are buried compared to the average proteome and some glycan structure. Anti-biotin and lectins are not drawn. N>30 for all measurements. All error bars indicate the standard deviation. p-values based on two-sample Student’s t-test are 0.54 and 0.0085.

In order for an immune cell to recognize and act on a tumor cell, like a HER2-positive breast cancer cell, it must be able to bind to a target molecule on the tumor cell surface, such as an antibody-bound antigen. To determine whether the measured proteome and glycome heights present a physical barrier to that binding, we measured the height of the therapeutic antibody target HER2 using the anti-HER2 antibody, Trastuzumab, the antibody-binding receptor FcγR, and the phosphatase CD45R_ABC_, which must be excluded for FcγR activation (Fig. 5a and b). We found the height of CD45R_ABC_ on Hoxb8 cells was 24.7 +/- 1.3 nm (S.D.), significantly exceeding the average total protein and glycan height, which would enable it to be excluded when the cells come into close contact. However, the measured heights of the therapeutic antibody Trastuzumab (7.9 +/- 1.4nm (S.D.)) and FcγR (6.2 +/- 1.7 nm (S.D.)) show that they are well below the average proteome heights on the native cell surface. This indicates that accessibility of HER2 and FcγR may be sterically limited by surrounding proteins. While these data are for cultured cells, our results suggest that variations in cell surface height profiles of tumor cells and immune cells in patients could significantly affect the accessibility of tumor antigens and the efficacy of antibody-mediated immunotherapies.

## Discussion

The cell surface landscape plays an important role in diverse cellular processes. Since small changes in the height of molecules can significantly alter receptor binding and glycoprotein exclusion from cell-cell interfaces, quantifying the height of molecules comprising the cell surface proteome and glycome with nanometer-scale resolution is important for a mechanistic understanding of cell-cell interactions.

We have developed a super-resolution localization-based method called Cell Surface Optical Profilometry (CSOP) that can rapidly and simply measure protein and glycan heights with nanometer resolution on native cell membranes. We applied this technique to measure membrane thickness, quantify the height and flexibility of purified as well as native proteins, and test the effect of surface and solution crowding on protein heights. Our all-optical approach can be multiplexed to simultaneously quantify the heights of multiple molecules in a matter of minutes with conventional spinning disk confocal microscopy, overcoming challenges with current methods that limit resolution, require extensive preparation time, or are not compatible with unfixed, native cell membranes.

CSOP takes the localization concept that enables 3D super-resolution imaging and trades lateral resolution for improved vertical resolution. Rather than the maximum practical resolution of ∼10 nm for PALM, STORM, and STED, CSOP is able to quantify average cell surface protein height with ∼1 nm resolution by averaging multiple molecules reporting the same heights. In addition to its precision, CSOP sample preparation is simple and the measurement is rapid. When measuring a native protein with commercially available antibodies, the cell preparation, surface labeling, calibration, measurement, and data analysis can be completed within 3-5 hours.

Though CSOP uses the straightforward concept of fluorophore localization, achieving nanometer-scale height resolution on native membranes required several developments to enable robust, high-precision measurements. First, we established a calibration protocol and membrane labeling strategy to localize membrane and proteins without errors associated with chromatic aberration or fluorescence polarization, allowing CSOP to maintain high resolution even with modest acoustic vibration (Fig. S8). Second, we validated the accuracy of average height measurement using double-stranded DNAs and MD simulations. Third, we quantified the offset introduced by antibody labeling, enabling height measurements and antibody epitope mapping of unmodified cell surface proteins, such as those on primary cells. Lastly, we confirmed that existing methods for preparing native cell membranes, including GPMVs and cell swelling, can be adapted to CSOP for the measurement of cell surface proteins on native cell membranes.

As shown above, CSOP has the potential to be a broadly useful tool for investigating diverse biological processes including protein isoform changes during cell differentiation (51), modifications in glycan height during cancer progression (26), and kinetics of protein conformation changes (52). Ongoing developments in protein labeling strategies using antibodies and site-specific chemical labels have the potential to improve the ability of CSOP to quantify the height and flexibility of cell surface molecules and their physiological impacts.

## Materials and Methods

### Materials

Size standard DNA oligonucleotides were purchased from Integrated DNA Technologies. Anti-biotin (BK-1/39; catalog number: sc-53179) was purchased from Santa Cruz Biotechnology. Monoclonal antibodies against mouse CD45 B220 (RA3-6B2; catalog number: 103201), CD45RB (C363-16A; catalog number: 103311), pan-CD45 (I3/2.3; catalog number: 147715, 30-F11; catalog number: 103123), and CEACAM5 (ASL-32; catalog number: 342302) were purchased from BioLegend. Antibody against Fc-receptor (2.4G2; catalog number: 553142) was purchased from BD Biosciences. Spot-Label ATTO488 (catalog number: eba488-10) was purchased from ChromoTek GmbH. Anti-SIRPα antibody (mSIRPα.20a) was kindly supplied by Aduro Biotech. The antibody mSIRPα.20a was derived from immunizing rats after several cycles of mouse SIRPα DNA immunization, followed by panning and selection of B cell clones producing antibodies that block the CD47 – SIRPα interaction. Clone 20a was found to cross-react to human (but not mouse) SIRPb1 and specifically bound to the N-terminal IgV domain. From the 20a Ig heavy and light chain sequences a mouse IgG1 chimera was constructed and produced in CHO cells for use in all experiments. Lysine-Cysteine-Lysine-Deca-histidine peptide was purchased from GenScript. Purified mouse CD45RABC extracellular domain with a C-terminal 6-His tag (accession #: NP_001104786, catalog number: 114-CD-050) was purchased from R&D systems. Lectins Sambucus Nigra Lectin (SNA; catalog number: L-1300), Maackia Amurensis Lectin II (MAL II; catalog number: L-1260), Soybean Agglutinin (SBA; catalog number: L-1010-10), and Wheat Germ Agglutinin (WGA; catalog number: L-1020-10) were purchased from Vector Laboratories. Fluorescently labeled cholera toxin subunit B (catalog number: C34775, C34776, C34778) was purchased from ThermoFisher Scientific. Latrunculin A (catalog number: ab144290) was purchased from Abcam. Y-27632 (catalog number: 1254) was purchased from Tocris. Paraformaldehyde (PFA; catalog-number: 16005) and Dithiothreitol (DTT; catalog number: D0632) was purchased from Sigma-Aldrich. Cell-Tak cell and tissue adhesive (catalog number: 354240) was purchased from Corning. 1,2-dioleoyl-sn-glycero-3-[(N-(5-amino-1-carboxypentyl)iminodiacetic acid)succinyl] (DGS-Ni-NTA; catalog number: 709404), 1,2-dioleoyl-sn-glycero-3-phosphocholine (DOPC; catalog number: 850375), 1,2-dioleoyl-sn-glycero-3-phospho-L-serine (DOPS; catalog number: 840035), 1,2-dimyristoyl-sn-glycero-3-phosphoethanolamine-N-(7-nitro-2-1,3-benzoxadiazol-4-yl) (14:0 NBD PE; catalog number: 810143), 1,2-dioleoyl-sn-glycero-3-phosphoethanolamine-N-[4-(p-maleimidophenyl)butyramide] (18:1 MPB PE; catalog number: 870012), 1,2-dimyristoleoyl-sn-glycero-3-phosphocholine (14:1 (Δ9-Cis) PC; catalog number: 850346). 1,2-dieicosenoyl-sn-glycero-3-phosphocholine (20:1 (Cis) PC; catalog number: 850396), 23-(dipyrrometheneboron difluoride)-24-norcholesterol (TopFluor Cholesterol; catalog number: 810255) were purchased from Avanti polar lipids. 1,2-Dioleoyl-sn-glycero-3-phosphoethanolamine labeled with Atto 488 (DOPE-488; catalog number: AD 488-16) was purchased from ATTO-TEC GmbH. Silica microspheres (2.47μm; catalog code: SS04N; lot number: 6911, 4.07μm; catalog code: SS05002; lot number: 12602, 5.04μm; catalog code: SS06N; lot number: 12477, 6.84μm; catalog code: SS06N; lot number: 4907) were purchased from Bangs laboratories.

### Cell culture and cell lines

HEK293T cells were obtained from UCSF Cell Culture Facility and grown in DMEM (Life Technologies) supplemented with 10% heat-inactivated FBS (Life Technologies) and 1% Pen-Strep (Life Technologies), at 37°C, 5% CO_2_. RBL-2H3 cells were obtained from UCSF Cell Culture Facility and grown in DMEM supplemented with 15% FBS and 1% Pen-Strep. HOXb8 cell is a generous gift from the Michael Sixt laboratory. It is grown in RPMI (Life Technologies) supplemented with 10% FBS, 1% Pen-Strep, 1μM β-estradiol (Sigma-Aldrich), and 100ng/ml FLT3 (PEPRO Tech). SK-BR-3 cells were purchased from ATCC (ATCC HTB-30) and grown in McCoy’s 5A (ATCC) supplemented with 10% FBS and 1% Pen-Strep. MDA-MB-453 cells were purchased from ATCC (ATCC HTB-131) and grown in L-15 (Life Technologies) supplemented with 10% FBS and 1% Pen-Strep without CO_2_. HEK293T cells were transiently transfected using TransIT-293 (Mirus Bio) to overexpress CEACAM5 containing an N-terminal SPOT-tag. RBL cell lines were generated using lentiviral transduction. To do this, CEACAM5 N-terminal domain, CEACAM5 full-length, SIRPα wild-type, SIRPα GPI-anchor, E-cadherin containing an N-terminal SPOT-tag were cloned into a pHR vector. The pHR construct was then co-transfected with second generation packaging plasmids, pMD2.G and p8.91, in HEK293T cells to generate lentivirus. RBL cells were transduced with lentivirus and used 24hours post transduction.

### Protein purification

Multi-FN3-domain proteins containing N-terminal ybbR-tags were expressed and purified as described previously (9). Briefly, FN3 proteins were expressed in Rosetta DE3 competent cells (EMD Millipore), lysed by sonication, and affinity purified over a His-Trap HP column (GE Healthcare). The proteins were gel-filtered and their size was validated via a Superdex 200 column on an AKTA Pure system (GE Healthcare). We labeled 50μM protein using 10μM SFP synthase (purified according to published protocols (53)), 40μM AlexaFluor dye-conjugated CoA, and 10mM MgCl_2_. The conjugates were then gel-filtered via a Superdex 75 10/300 GL column. The extracellular human CEACAM5 N-terminal domain and full-length proteins (Uniprot P06731), mouse SIRPα (Uniprot P97797), and mouse E-cadherin (Uniprot P09803) were purified as described previously (9). Briefly, HEK293T cells were transfected with a construct consisting of CEACAM5 protein, SIRPα, or E-cadherin with an N-terminal ybbR-tag and a C-terminal His-tag in a pCAGGS vector at 70% confluency. After 48 hours, the supernatant was collected for purification using His Trap Excel column (GE Healthcare). The proteins were labeled with an AlexaFluor dye through the ybbR-tag and gel-filtered via a Superdex 75 10/300 GL column, selecting for only labeled proteins.

### Microscope for CSOP height measurements

CSOP imaging was carried out on an inverted Nikon Eclipse Ti microscope (Nikon Instruments) equipped with a Yokogawa CSU-X spinning disk using an oil-immersion objective (Apo TIRF 100x, numerical aperture (NA) 1.49, oil). Three solid state lasers were used for excitation: 488nm, 561nm, and 640nm (ILE-400 multi-mode fiber with BCU, Andor Technologies). The laser power at the sample plane was less than 1.5mW for all three channels. Fluorescent light was spectrally filtered with emission filters (535/40m, 610/75, and 665LP, Chroma Technology) and imaged on a sCMOS camera (Zyla 4.2, Andor Technologies). Z-stack was acquired using a piezo z-stage (npoint).

### Calibration of CSOP height measurements to account for chromatic aberrations

Prior to making CSOP measurements, a calibration procedure was run once for a microscope configuration (see Fig. S2). The calibration begins with taking a z-stack of a lipid-coated glass bead with the two fluorophores to be used for the height measurement both labeling the same lipid bilayer, which should give a height measurement of zero. Based on the images, we calculate the radius of fluorophores in each z-slice using a custom MATLAB script (see Protocol for CSOP data analysis) and determine the maximum radius in each color at the bead equator. The offset between the measured fluorescent radii (h_offset_) results solely from the chromatic aberration, providing a parameter that can be used to compensate subsequent measurements. When the measured height, h_measured_, of a cell surface molecule is obtained by quantifying the difference in radii of the fluorescently-labeled bilayer and tip of the surface molecule, the aberration corrected average height, <h>, can be calculated as <h> = <h_measured_> - <h_offset_>.

### Method for avoiding polarization effects

Separate from the chromatic aberration, polarization effects of fluorophores labeling lipid membranes can also induce a non-zero <h_offset_> for beads of diameter 8 μm or less. The majority of lipid-fluorophore conjugates are highly oriented by the bilayer leading to errors in the correction of cell surface molecular height measurements (32). To avoid this, we used an AlexaFluor dye-conjugated to a His-tag that binds to the surface of Ni-containing bilayers in place of lipid-fluorophore conjugates to localize the bilayer surface (Fig. S3).

### Protocol for CSOP height measurements

CSOP sample preparations are described in the sections below. The samples were loaded on a custom imaging chamber, which was made by adhering a PDMS well to a 1.5 coverslip (Thermo Fisher Scientific). The coverslip was passivated by forming a SLB or by adsorbing BSA while presenting binding molecules such as Cell-Tak for immobilizing CSOP samples. Prior to CSOP imaging, the laser power and exposure time was adjusted in both channels. In general, a <1 mW laser power at the sample plane and an exposure time shorter than 200ms provided a SNR ∼10 or higher in both fluorescence channels. We kept the SNR of two fluorescence channels identical to avoid any potential bias to CSOP measurement. We manually determined the equator of samples and acquired approximately 20 slices of images with a 50 - 100 nm step size using the z-piezo stage. We collected images in two fluorescence channels before moving the z position, but collecting the z-stack of each fluorescence channel separately did not affect the results. Each z-stack image contained up to ten spherical samples per field-of-view.

### Protocol for CSOP data analysis

We used a custom MATLAB (Mathworks) script to estimate the radius of samples from CSOP images (Supplementary Software). The algorithm radially averages fluorescent signals from the center of a circle to construct the 1D signal distribution in the radial direction. To determine the center of a circle accurately, we performed the radial averaging in 900 different center locations and quantified the height of radial profile at its peak. We then created the 2D map of peak heights versus centers and fit the 2D Gaussian surface to estimate the true center, where the peak height becomes maximized. To determine the peak location of the radial profile from the center, which corresponds to the radius of CSOP sample, we fit the Gaussian curve to the top 10% region of the peak.

### Measurement of lipid bilayer thickness

To create a lipid-coated bead, first small unilamellar vesicles (SUV) were formed according to published protocols (9). To calibrate the measurement baseline, a lipid film composed of 20:1PC, 2.5 mol% Ni-NTA, 0.15mol% DOPE-LissRhod B, and 1 mol% TopFluor-Cholesterol was rehydrated with MQ water, sonicated at low power (20% of max), added to MOPS buffer (final concentration, 50mM MOPS pH 7.4, 100mM NaCl), and filtered through a 200nm PTFE filter (Millipore). Next, 10uL of 6.8µm glass bead slurry (10% solid) were cleaned using a 3:2 mixture of H2SO4:H2O2 (30 minutes in bath sonicator) and cleaned three times in MQ water. Finally, 5µL of clean bead slurry were added to 50µL of SUV solution and mixed gently by pipetting. The bead/SUV mixture was incubated for 15 minutes at room temperature before the mixture was washed with HEPES buffer (50mM HEPES pH 7.4, 100mM NaCl) for three times by gentle syphoning. CSOP measurement was performed to determine the offset between bilayer mid-planes measured by TopFuor versus LissRhod B. To measure bilayer thickness, lipid-coated beads containing only Liss Rhod B was formed. The lipid-coated beads were subsequently incubated with TopFluor-cholesterol in solution, which spontaneously incorporates into the bilayer’s outer leaflet. 25µM of TopFluor-Cholesterol was first mixed with 50µM Methyl-β-cyclodextrin (MβCD) to make the cholesterol compound soluble in buffer. By comparing the locations of TopFluor-cholesterol (mid-plane of bilayer) versus LissRhod B (surface of bilayer), we determined bilayer leaflet thickness. We performed the measurement for the lipid composed of 14:1PC as well as 20:1PC.

### Measurement of DNA length

6.8μm lipid-coated beads were formed using lipids composed of DOPC, 3.5 mol% Ni-NTA, and 14:0-NBD PE. The beads were incubated with the size standard DNAs (Table S2) containing an Atto555 dye on one end and cholesterol on the other in buffer containing 150 mM HEPES and 300mM NaCl for 10 minutes before imaging. We added varying amounts of DNA to obtain the same surface density between DNA standards. For example, 1 nM of 15 bp or 50 nM of 105 bp provided comparable surface density of ∼1000/μm^2^. The location of membrane surfaces was measured by binding AlexaFluor 555-conjugated deca-histidine peptides to the Ni-NTA lipids.

### Numerical simulation of a double-stranded DNA height

We assumed a dsDNA as a worm-like chain with variable persistence length and 0.35 nm per basepair. We determined the probability distribution of its end-to-end distance based on an analytical expression previously derived by Murphy and Ha (54). To determine the average height of a DNA while its one end is anchored on a surface and the other end is freely rotating in 3D, we assumed a hemispherical volume where its internal density distribution in radial distance is equal to the probability distribution of DNA end-to-end distance. We then determined DNA height by calculating the center-of-mass of the hemisphere. We calculated the deviation of CSOP measurement from the simulated DNA height at varying persistence length and found that the error is minimized when the persistence length is approximately 50 nm.

### Measurement of multi-domain protein height and crowding effects

6.8μm lipid-coated beads were formed using lipids composed of DOPC, 3.5 mol% Ni-NTA, and 14:0-NBD PE. The beads were incubated with 10nM of FN3 single- or multi-domain protein containing an AlexaFluor555 dye at its N-terminus and a H10-kck-H6 tag at its C-terminus in HEPES buffer (50mM HEPES and 100mM NaCl, pH 7.4) for 20 minutes before imaging. The surface density of proteins was approximately 500/μm^2^ for all sizes measured. The location of membrane surfaces was measured by binding AlexaFluor 555-conjugated deca-histidine peptides to Ni-NTA lipids. To test the effect of protein surface crowding on height, we bound 11XFN3 proteins at 10nM, 40nM, and 180nM, which provided surface density of ∼460/µm^2^, 1380/µm^2^, and ∼3470/µm^2^. To test the effect of PEG in solution on protein height, we made lipid-coated beads using lipids composed of DOPC, 5 mol% DOPS, 5 mol% 18:1 MPB-PE, and 14:0-NBD PE. We then covalently bound 9XFN3 proteins to the bilayer using their C-terminal cysteines by incubation for 1 hour at room temperature. This was important, because proteins bound via histidine-Nickel interactions became unbound quickly after PEG addition. Following protein binding, PEG8k was added to solution to a final concentration of 2.5%, 5%, 10%, or 20% w/v before imaging.

### Measurement of multi-domain protein height on GPMVs

RBL cells over-expressing a recombinant protein containing an N-terminal SPOT-tag were created by lentiviral transduction. Giant plasma-membrane vesicles (GPMV) were formed by adding 25 mM PFA and 2mM DTT to cells (44). GPMVs were collected within 1 hour and subsequently incubated with SPOT V_HH_ conjugated with Atto488 and cholera toxin subunit B-AlexaFluor555 for 20 minutes at room temperature before imaging. GPMVs remained taut throughout imaging. The location of membrane surfaces was measured by the cholera toxin subunit B-AlexaFluor488 and 555 pair.

### Measurement of multi-domain protein height on swollen cells

HEK293T, HOXb8, SKBR3, and MDA-MB-453 cells were used in swelling experiments. Cells were lifted by shear and transferred to an imaging chamber coated with Cell-Tak cell adhesive according to the manufacturer protocol. Cells were loaded at a density where single cells were well separated and then allowed to gently adhere to the glass for 10 minutes at room temperature. Cells were subsequently washed once with PBS, and cytoskeletal drugs were added (Latrunculin A and Y27632 at 1μM and 10μM) for 30 minutes at room temperature or 37°C. Cells were washed three times with PBS, and labeling reagents, such as a labeled antibodies, lectins, or CellMask as well as cytoskeletal drugs, were then added for 20 minutes at 4°C. HEPES-buffered MQ (pH 7.5) containing cytoskeletal drugs was subsequently added to swell cells. MQ was added in several steps while cells were monitored. Typically, cells were fully swollen in between 1/4th and 1/10th PBS and remained taut for at least one hour.

### Simulation of CSOP height measurement resolution

We numerically simulated CSOP images to test for CSOP resolution and its sensitivity to acoustic vibration using a custom Matlab script (Supplementary Software). To do this, we generated an image of a circle reflecting an CSOP sample image at the equatorial plane using the 2D Gaussian point spread function with a standard deviation of 130 nm, a camera pixel size of 55 nm, and Poisson shot-noise. We first generated a true image of a circle and convoluted the circle with the PSF. We then added Poisson noise in each pixel by using the original signal as the mean. We created a pair of circle images simulating the membrane and the protein and determined the protein height by estimating the radii of each circle. We repeated this for 25 pairs of images that were identical except for photon-shot noise and used the root-mean-square-error of protein height as an error (Table S1).

### Effect of vibrations on CSOP height measurements

To simulate acoustic vibration (Fig. S8), we allowed a circle as described in the previous section to move 5 steps in a random orientation and accumulated images of circles in 5 different positions to make one image frame.

### Determination of surfaceome amino acid length distribution

We used human surfaceome database compiled by the Wollscheid group (2), where protein domain and topology are annotated for approximately 3000 proteins comprising human surfaceome. Amino acid length of protein extracellular domain was obtained by counting the entire extracellular domain in the case of GPI-anchored and single-pass membrane proteins or by counting the longest extracellular domain in the case of multi-pass membrane proteins. Extracellular domain length was variable from 5 to 22096 amino acids for proteins, while 51% of proteins contain more than 150 amino acids in their extracellular domain. These proteins are likely to extend taller from the cell surface when compared to a single Ig-domain, which contains approximately 110 amino acids.

### MD simulation overview

To explore a molecular model of protein dynamics on fluid membranes, we performed coarse-grained molecular dynamics (MD) simulations of semi-flexible polymers diffusing on 2D surfaces. We model protein chains using a standard Kremer-Grest bead-spring model (55), with each bead representing a structured protein domain (Fig. S4). A bead at one end of the chain is confined between two parallel walls separated by one bead diameter, allowing it to diffuse freely in 2D but cannot escape out of plane. All other beads on the chain are free to move in 3D, except through a bottom wall that acts as a solid substrate, thereby modeling protein diffusion along a membrane that is in-plane fluid and out-of-plane elastic. Because the size of each protein domain is large compared to the surrounding solvent molecules, the solvent is coarse-grained out and its dynamics are not explicitly modeled. In other words, the protein chain experiences a hydrodynamic drag and Brownian motion from the continuous solvent. In this work, membrane thermal fluctuation was not considered.

Simulations were performed using a GPU-enabled HOOMD-blue molecular dynamics package (56, 57), and all simulation results constitute an average of over at least 2000 protein chains. The dynamics of particle bead *i* is evolved in time following the overdamped Langevin equation

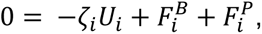

where *ζ_i_* is the hydrodynamic drag factor, *U*_*i*_ is the velocity, 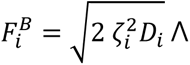 is the translational Brownian force, *D*_*i*_ is the Stokes-Einstein-Sutherland translational diffusivity of a single bead, and 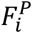 is the interparticle force. The left-hand side is zero since inertia is negligible for coarse-grained proteins embedded in a viscous solvent. The translational diffusivity is modeled with the usual white noise statistics, 〈Λ(t)〉 = 0 and, 〈Λ(t)Λ(0)〉 = *δ*(*t*), where *δ*(*t*) is a delta function and *I* is the identity tensor. The drag coefficient *ζ*_*i*_ of particle *i* is linearly scaled with the particle size *σ*_*i*_. The inter-particle force (described below) includes contributions from intra-chain potential, inter-chain bead-bead pair interactions, and bead-wall interactions.

All bead pair interactions and bead-wall interactions are modeled with a Weeks-Chandler-Andersen (WCA) potential (58), in which a Lennard-Jones (LJ) potential is shifted upwards, truncated at the potential minimum of 2^1/6σ^ (such that the potential is purely repulsive), and assigned a well depth of *∈* = *k*_*B*_*T* (where *k*_*B*_*T* is the thermal energy). All lengths are expressed in units of the LJ diameter *σ* and are set to unity. We connect the polymer chains with a finitely extensible nonlinear elastic (FENE) potential, using a spring constant of *k* = 30 and a bond length of *R*_0_ = 1.5 (expressed in terms of reduced LJ units *ε* = *σ* = 1).

We used a system box size of *V* = L^2^L_z_ with L_z_ = 50*σ* and L was adjusted to achieve the specified area density and number of polymer chains (Fig. S4). Dilute surface densities were conducted with ∼200 chains/μm^2^, and the dense surfaces went up to ∼20000 chains/μm^2^. We imposed periodic boundary conditions in all directions, although the direction tangential to the substrate does not require them because the bottom wall prevents any particles from passing through. Initial configurations are generated by placing the particles in lattice locations, and sufficient time steps are run to reach a steady-state. Time steps were varied from Δ*t* = 10^−6^ − 10^−4^ and verified to be sufficiently small to capture relevant dynamics.

### MD simulation: Cell surface molecules as semi-flexible polymers

To model semi-flexible polymers, we implemented a bending potential between 3 neighboring particles to capture chain stiffness, *U*_B_ = *∈*_B_(1 − cos(*θ*_ijk_ − *θ*_0_)), where *∈*_B_ is the bending energy, *θ*_ijk_ is the bond angle between neighboring particles (i, j, k), and *θ*_0_ = *π* is the resting angle (Fig. S4 and Movie S1). The persistence length can be measured by calculating the bond angle correlation, 〈*e*_1_ ⋅ *e*_i_〉 = exp(−*s*_i_/*ℓ_P_L_p_*), where *e*_i_ is the unit vector connecting the center of mass positions of particles i and i+1, and *s*_i_ is the path length along the polymer to particle i. To implement the persistence length *L_p_* as an input to the simulations, the bending energy was simply set to *∈*_B_/(*k*_B_*T*) = *ℓ_P_L_p_*/*σ*. The bond angle correlation of an unbound protein exhibited an exponential decay, validating our implementation of persistence length via bending stiffness (Fig. S5). Interestingly, we observed that, for a polymer bound on a wall, the bottom wall locally stiffens the polymer due to its interactions with the wall, elevating the polymer height compared to an unbound polymer (Fig. S6).

### MD simulation: Solution effects on cell surface molecular height

For simulations of macromolecular osmotic compression, spherical particles were added to the bulk of the simulation box to model the effects of macromolecular crowding. To model PEG8k used in the experiments, these bulk particles were chosen to have a soft repulsive potential between particle pairs. This was implemented in HOOMD by using a DPD conservative pair force between all pair interactions involving the bulk PEG particles, with the force coefficient *A* = 25.0 and a cut-off distance of 2^1/6^*σ*_ij_, where *σ*_ij_ is the distance between particle pairs i and j. The radius of gyration of PEG8k was estimated to be within the range of 2.5-3.4 nm (59–61), which forms the upper and lower bounds of simulation data (Fig. 2e and Movie S2).

It is an interesting outcome that the sole effect of PEG was to compress the proteins. One may think that the PEG may also cause pairs of protein chains to aggregate together through a depletion flocculation mechanism. We did not observe any large-scale protein clustering, presumably because the bottom substrate is an even better depletion surface for a protein chain and because we are at low surface densities of proteins in experiments.

To study the effects of macromolecular osmotic compression on a mixture of proteins of different sizes, we conducted simulations with 5% w/v PEG8k in the bulk and an equal mixture of surface proteins with lengths 3*σ*, 9*σ*, and 20*σ* at three different total concentrations: dilute 200 chains/μm^2^, intermediate 5000 chains/μm^2^, and dense 20000 chains/μm^2^. At all three total densities, the absolute magnitude of compression was largest for the tallest protein (20*σ*). At 20000 chains/μm^2^, the protein heights decrease from 8.2 nm, 22.2 nm, and 46.0 nm without any added PEG to 6.1 nm, 19.2 nm, and 42.4 nm with 5% w/v PEG8k for the 3*σ*, 9*σ*, and 20*σ* proteins, respectively.

### MD simulation: Height offset added by antibody labeling of cell surface molecules

For simulations of antibody labeling of cell surface proteins, we constructed an antibody using a rigid assembly of 5 spheres using physical dimensions of IgG antibodies: ∼15 nm wide, ∼8 nm tall, and ∼4 nm thick (62, 63) (Fig. S7 and Movie S3). We assign one of the Fab domains to bind strongly to a single domain on the protein chain. The attractive pair potential between these two particles is radially symmetric such that the antibody can swing freely in all angles around the protein binding site. All other particle pairs experience a short-ranged repulsive WCA potential, as described earlier. We initialize the simulation with the antibody binding to the protein, and after sufficient equilibration, we calculate the average offset of the center of mass of the antibody relative to the binding site on the protein. Although it is known that antibodies are flexible macromolecules (63, 64), we modeled them as a rigid assembly for simplicity. Flexibility is straightforward to include in the model, but because we are calculating the average offset of the center of mass of the antibody relative to the protein, we do not anticipate antibody flexibility to influence our results.

We find that the average height of the center of mass of the antibody is always larger than its binding site on the protein chain. This is a result of steric interactions between antibody-substrate and antibody-protein pairs. As the protein chain and antibody undergo Brownian fluctuations, the Fab and Fc regions of the antibody experience short-ranged repulsive potentials when interacting with the protein and substrate. This causes a positive average height difference between the antibody and its binding site. For antibodies binding to short proteins, the offset is larger because the substrate constrains the antibody to remain above the surface. As the protein length increases (and hence the antibody binding site also increases in height), the average offset decreases because the antibody can explore more configurations below the binding site. The offset remains positive because steric interactions with the protein constrain the antibody to explore regions above the protein on average.

To quantify this height offset in more detail, we conducted a series of simulations. First, we varied the overall length of the protein at dilute surface densities to measure the offset as a function of the average height of the protein terminal domain. We conducted this simulation at persistence lengths L_*p*_ = 12 nm and L_*p*_ = 0 nm and find that both curves are quantitatively similar, indicating that protein flexibility does not contribute significantly to the antibody-protein height offset under these conditions. We obtained similar results with L_*p*_ = 12 nm at larger surface density of proteins, 10000 chains/μm^2^, indicating that protein crowding also does not contribute largely to antibody height offsets. Lastly, we fixed the overall protein length to 10L (with average terminal domain height ∼ 15.5 nm), and varied the position of the antibody binding site along the protein length. Because of the symmetry of steric interactions between antibody-protein pairs above and below the binding site, the offset is slightly smaller but remains quantitatively similar to the rest of the conditions. Because the width of the antibody is ∼15 nm, its interaction with the substrate plays a large role regardless of where it binds to the protein. However, for a very tall protein with average height 40 nm and antibody binding at sites higher than 20 nm, the offset becomes appreciably smaller than the antibody binding to terminal protein domains at equivalent heights. For example, an antibody binding at a 20 nm height has an offset that decreases from ∼1.5 nm to ∼0.5 nm when the protein has a full length of 40 nm (i.e., antibody binds at the middle of the protein).

All of these simulations taken together reveal that for most cell surface proteins of average height smaller than 15 nm, the offset of the antibody above its binding site is quantitatively similar regardless of protein flexibility, surface density, and antibody binding position along the protein. This prediction is corroborated by experimental data as shown in the main text.

## Acknowledgments

The authors would like to thank the Fletcher Lab members for feedback. The authors thank Dr. Erik Voets and Aduro Biotech for supplying the anti-SIRPα antibody. The illustration of proteins in Figure 2c are made using cellscape by Jordi Silvestre-Ryan (https://github.com/jordisr/cellscape). This work was supported by the NIH R01 GM114671 (DAF), the Immunotherapeutics and Vaccine Research Initiative at UC Berkeley (DAF), the Miller Institute for Basic Research, and the Chan Zuckerberg Biohub (DAF). S.S. was funded by a LSRF fellowship. B.B. was supported by the NIH Ruth L. Kirschstein NRSA fellowship NIH (1F32GM115091). S.C.T. acknowledge support from the Miller Institute for Basic Research in Science at U.C. Berkeley. D.A.F. is a Chan Zuckerberg Biohub investigator.

## Author Contributions

S.S. conceived of the study, designed and performed the experiments, analyzed the data and wrote the manuscript. S.C.T performed MD simulations and wrote the manuscript. B.B. designed experiments and interpreted the data. M.P. designed and performed the experiments. M.H.B. conceived of the study and analyzed data. D.A.F. conceived of, designed and supervised the study, interpreted the data and wrote the manuscript. All authors reviewed and approved the manuscript.

## Supplementary Information Appendix for

**Fig. S1:**
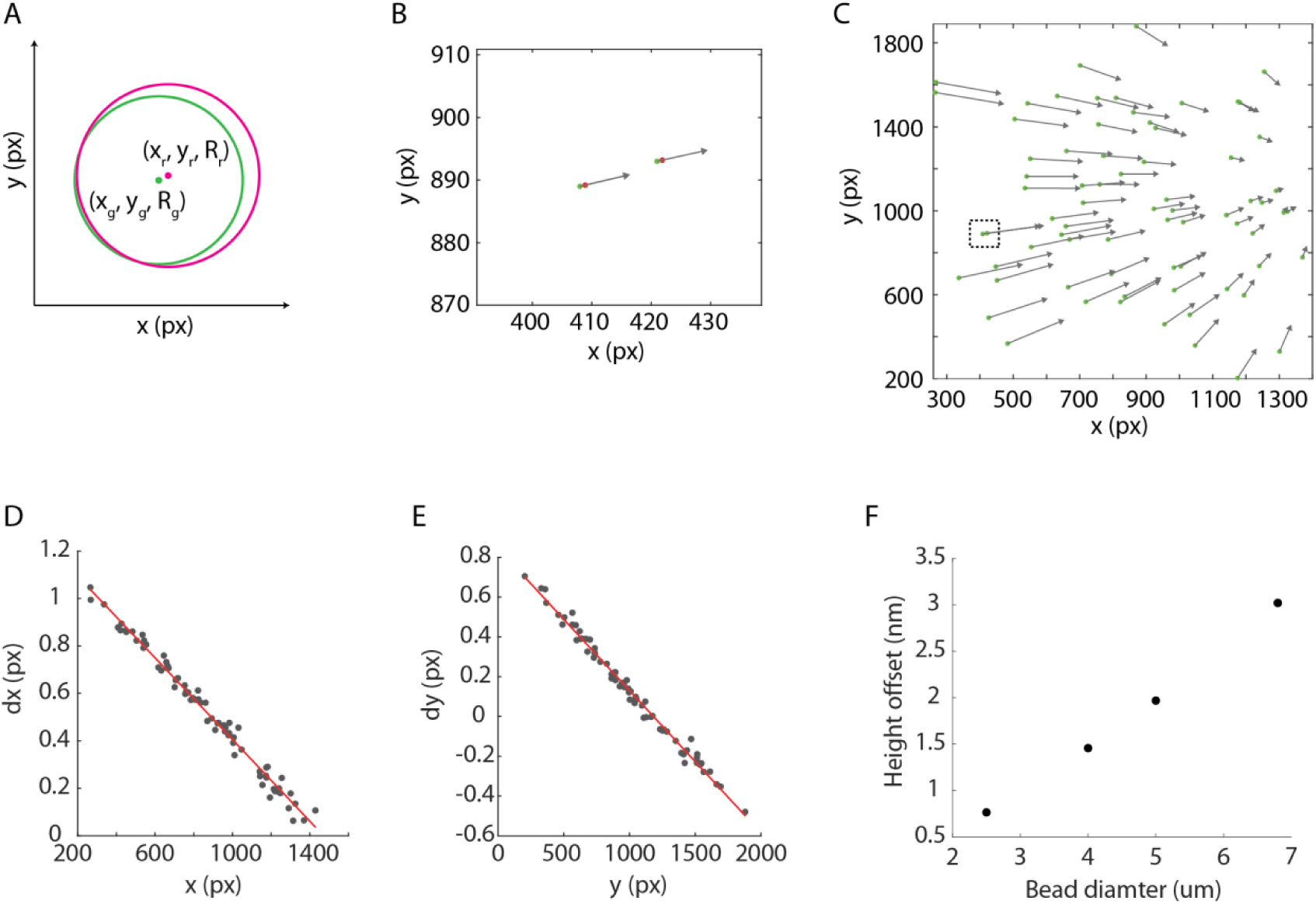
Characterization of CSOP chromatic aberration. (A) Schematic of chromatic aberration in 2D. The transformation between two-color channels is typically constructed by comparing the location of a single fluorophore in two different color channels. To perform this in CSOP, we compared the center of a lipid-coated bead imaged in two different color channels. (B) The centers of two representative beads – lipid-coated beads of size 5 µm – were estimated in green and red fluorescence channels (green and red dots). The grey arrows indicate vectors connecting the green and red centers. The magnitude of a vector is not drawn to scale. (C) The transformation vectors between the green and red channels for CSOP measurements on our microscope are constructed by measuring approximately 100 lipid-coated beads. The dashed grey box is zoomed in (B). (D and E) The scaling of a circle between the green and red channels (or any pair of fluorescent channels) can be calculated using the transformation vectors in C. In this example, the scaling dx/x and dy/y is 0.00078 and it is linear in both x and y (R-square of the fit is 0.986 and 0.988). This indicates that the chromatic aberration-induced height offset in a 5 µm bead is 1.9 nm. (F) Chromatic aberration-induced height offset for various bead sizes.

**Fig. S2:**
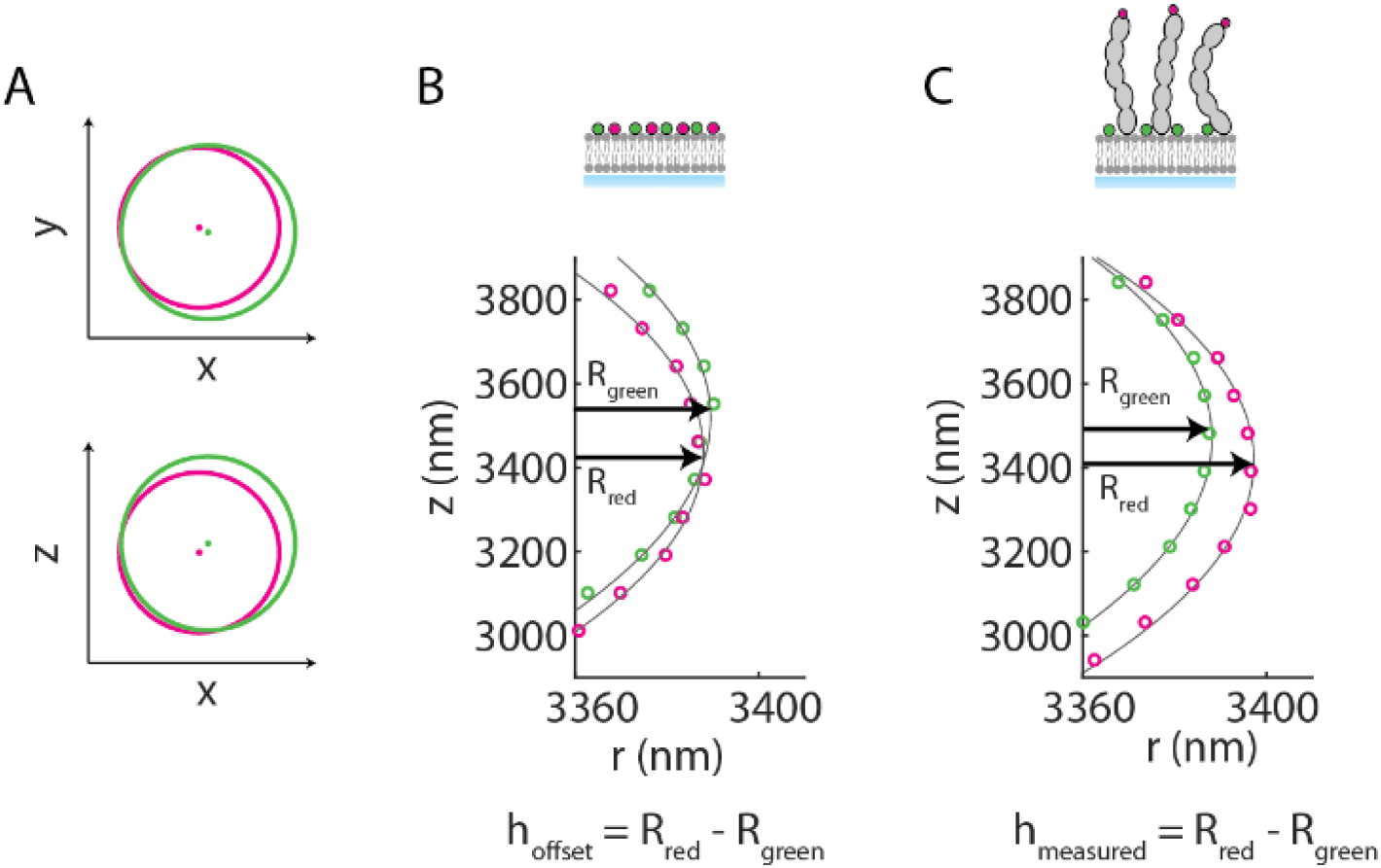
Calibration of CSOP height measurements. (A) Schematic illustrations of the effect of chromatic aberrations on the image of a lipid-coated bead labeled in green and red. The images between two channels are shifted and scaled in both x-y (lateral) and x-z (axial) planes due to aberrations. (B) h_offset_ calculation. For a lipid-coated bead labeled with green and red dyes on the bilayer surface, radius measured at different z locations (the green and red circles) were fit using a circle function (the grey lines) and the equator radius R_red_ and R_green_ were determined. The offset between the two radii, h_offset_, originates from chromatic aberration. (C) Calculation of h_measured_. For labeled proteins bound to a lipid-coated bead, the average difference between the equator radii, R_red_ and R_green_, <h_measured_>, is the protein height, <h>, shifted by <h_offset_>.

**Fig. S3:**
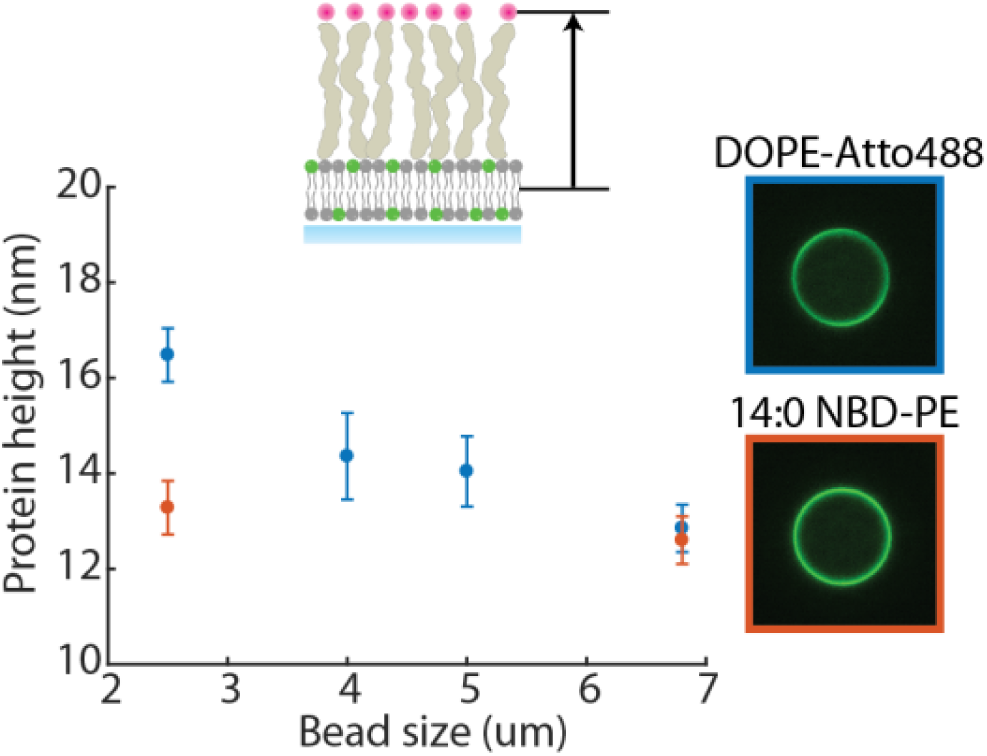
Effect of fluorescence polarization on CSOP height measurements. FN3 6L protein is measured on a lipid coated-bead where lipids are labeled with either DOPE-atto488 dye or 14:0 NBD-PE dye. DOPE-atto488 is significantly polarized in the bilayer as depicted in the micrograph, while 14:0 NBD-PE is polarized less. Height measured with a DOPE-atto488 containing bead (blue) shows a positive offset from the height measured with a 14:0 NBD-PE containing bead (orange). The offset, which is more than 3 nm for a 2.5 µm bead, decreases as the bead size increases. The error bar is the standard deviation.

**Fig. S4:**
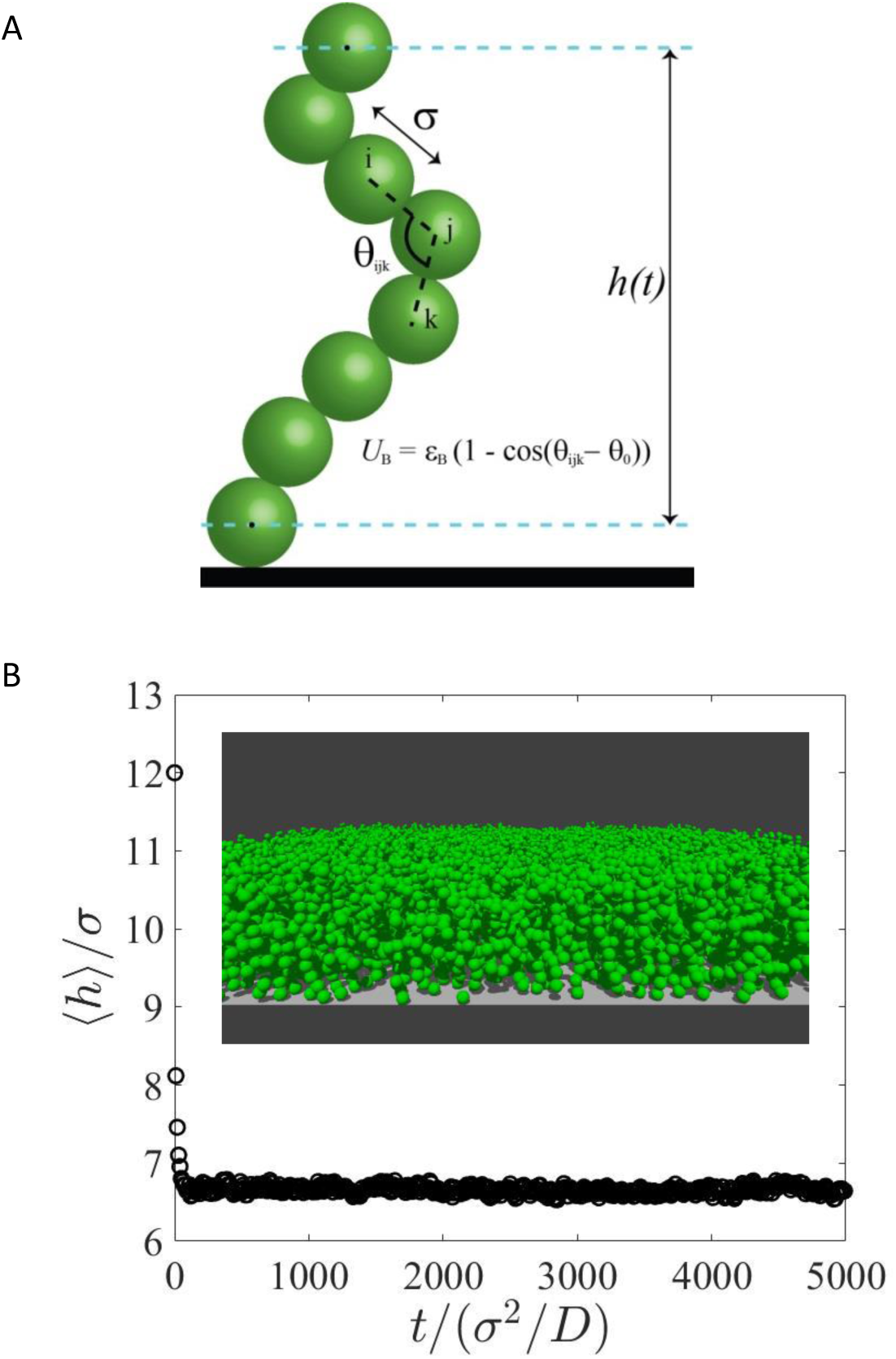
MD simulations of multi-domain proteins. (A) Schematic of a single polymer protein chain modeled in MD simulations. Using a Kremer-Grest bead-spring model, individual beads of diameter σ are connected by a FENE potential and a bending stiffness is invoked by a 3-particle angular potential, *U*_B_. The bottom bead is confined to remain along a 2D surface. The height of the center of mass of the top bead is recorded in time. (B) Simulation measurement of the height of the center of mass of the top bead relative to the substrate. Height is non-dimensionalized with bead diameter *σ* and time with *σ*^2^/*D*, where *D* is the Stokes-Einstein-Sutherland diffusivity of a single bead. Each datum is an ensemble average of over 2000 protein chains. The simulation is initialized with all proteins standing vertically upwards and quickly relaxing to a steady-state height. Results here are shown for a chain of 11 beads at a surface density of ∼15000 chains/μm^2^. The inset shows the snapshot of simulation.

**Fig. S5:**
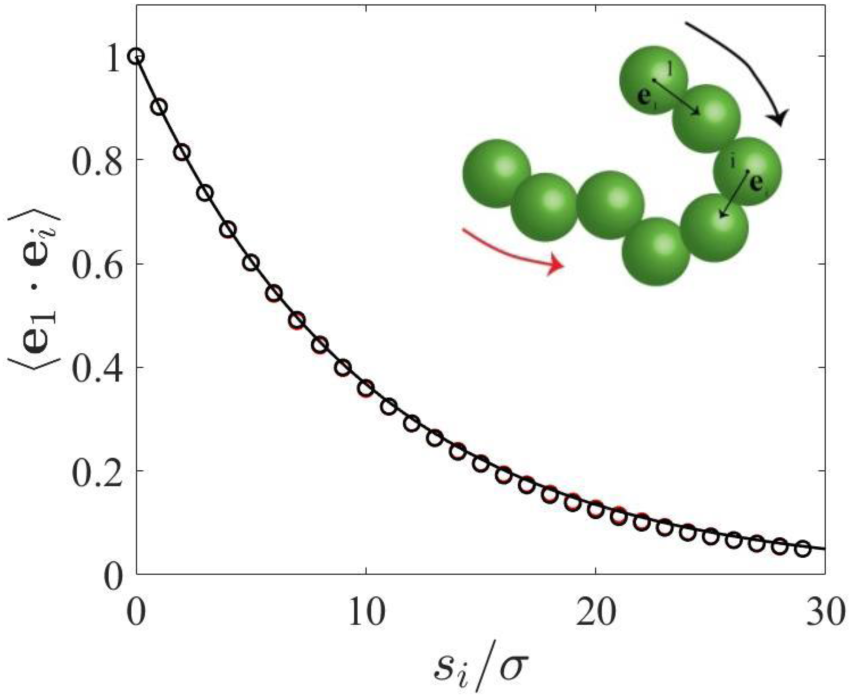
MD simulations of multi-domain proteins as semi-flexible polymers. Persistence length measurement of an unbound protein via the bond angle correlation (in black circles), where *e*_i_ is the unit vector connecting the center of mass positions of particle i and i+1, and *s*_*i*_ is the path length along the protein to particle i. The black solid curve is the theoretical prediction, 〈*e*_1_ ⋅ *e*_i_〉 = exp(−*s*_i_/L_*p*_), for a given input persistence length, L_p_. Data shown are for a chain of contour length of 30*σ* and L_*p*_ = 10*σ*.

**Fig. S6:**
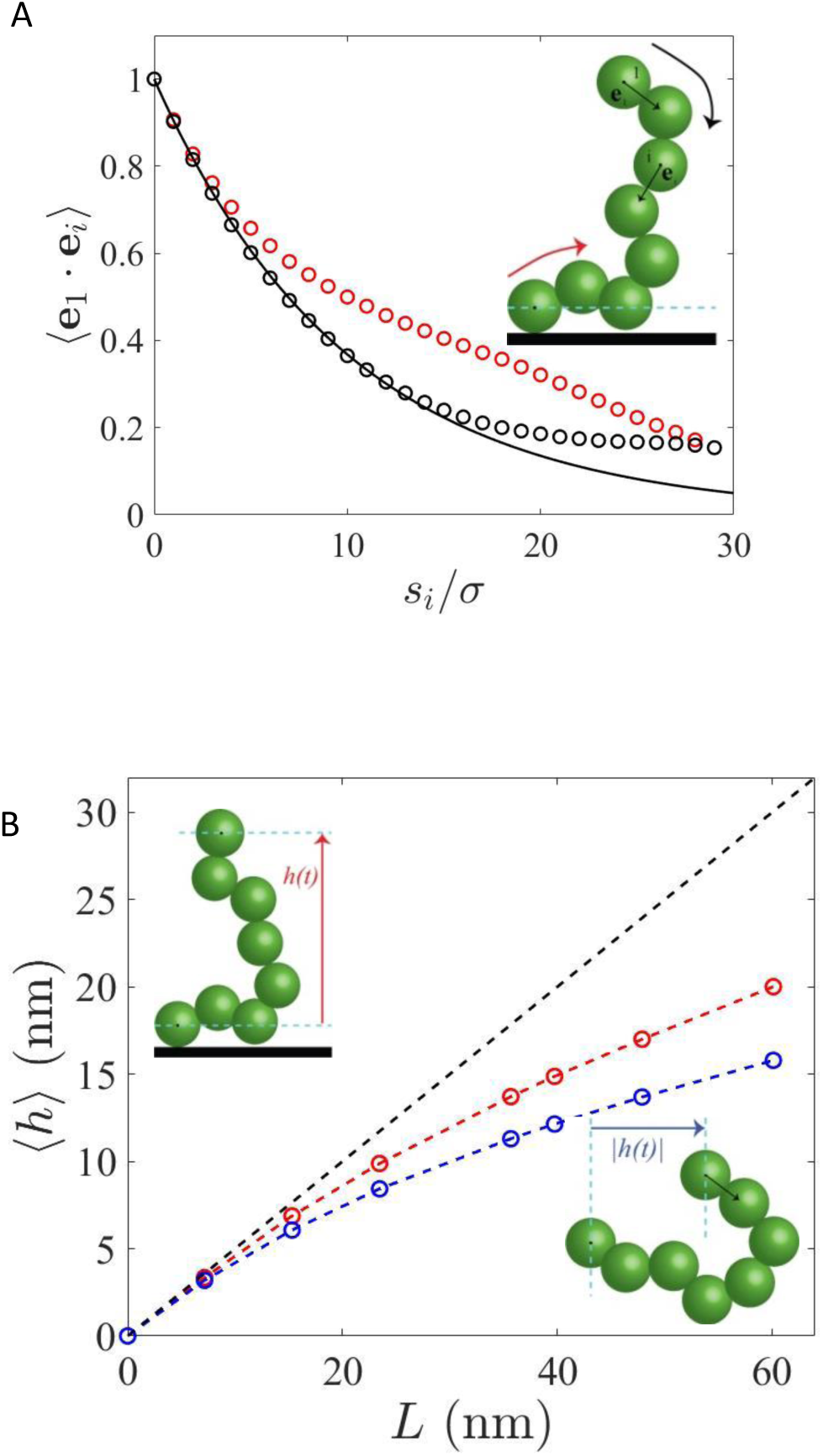
MD simulation of multi-domain protein height on a surface. (A) Correlations between particle vectors were calculated as in Fig. S5, from two directions of the polymer: from the wall outwards (in red circles) and from the bulk towards the wall (in black circles). A protein bound on a surface exhibits different relaxation of 〈*e*_1_ ⋅ *e*_i_〉 depending on the direction of measurement, demonstrating that the polymer is locally stiff near the wall. (B) Average height of a protein with one end bound to a surface (top curve, in red) and an unbound protein in 3D (bottom curve, in blue). For the unbound protein, the magnitude of the vertical distance between the first and last beads are reported, effectively “reflecting” the heights to the positive plane.

**Fig. S7:**
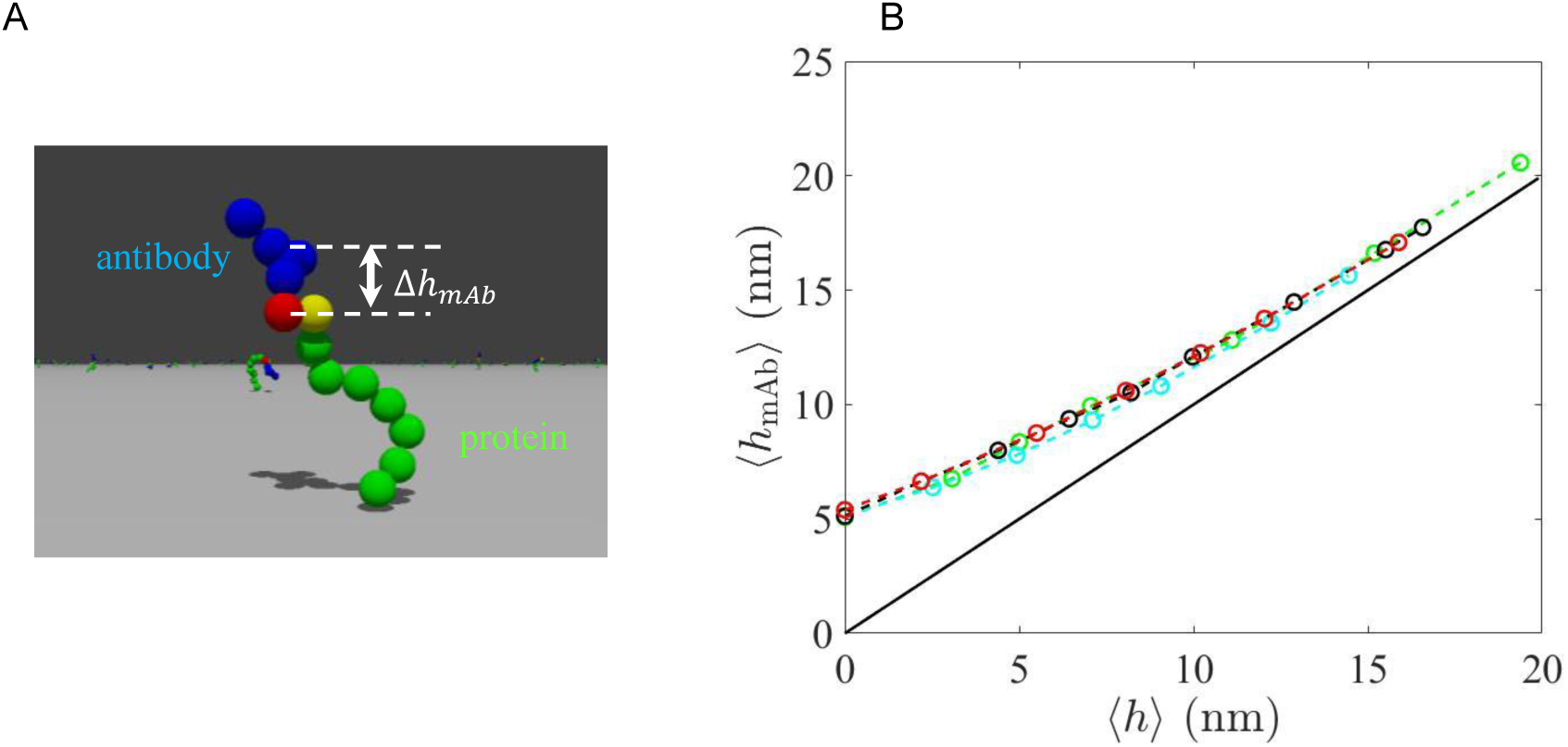
MD simulation of the height offset added by an antibody. (A) Simulation snapshot of an antibody binding to the terminal domain of an 8-domain protein. The antibody is modeled as a rigid assembly of 5 spheres using typical dimensions of an IgG antibody. The red particle represents a Fab region on the antibody that binds to the terminal domain of the protein (shown as a yellow particle). The offset of the center of mass of the antibody relative to its binding site on the protein is measured in the simulations. (B) Average height of the center of mass of the antibody as a function of the average height of its binding site on the protein. Four different sets of simulations were conducted: antibody binding to the terminal domain of proteins of various lengths with persistence length *L_p_* = 12 nm at dilute surface density (black symbols); same conditions but with *L_p_* = 0 nm (green symbols); same conditions with *L_p_* = 12 nm but at dense surface density of 10000 chains / μm^2^ (red symbols); lastly, antibody binding to various domains of a 10L-length protein (with average tip height ≈ 15.5 nm), with *L_p_* = 12 nm at dilute surface density (cyan symbols).

**Fig. S8:**
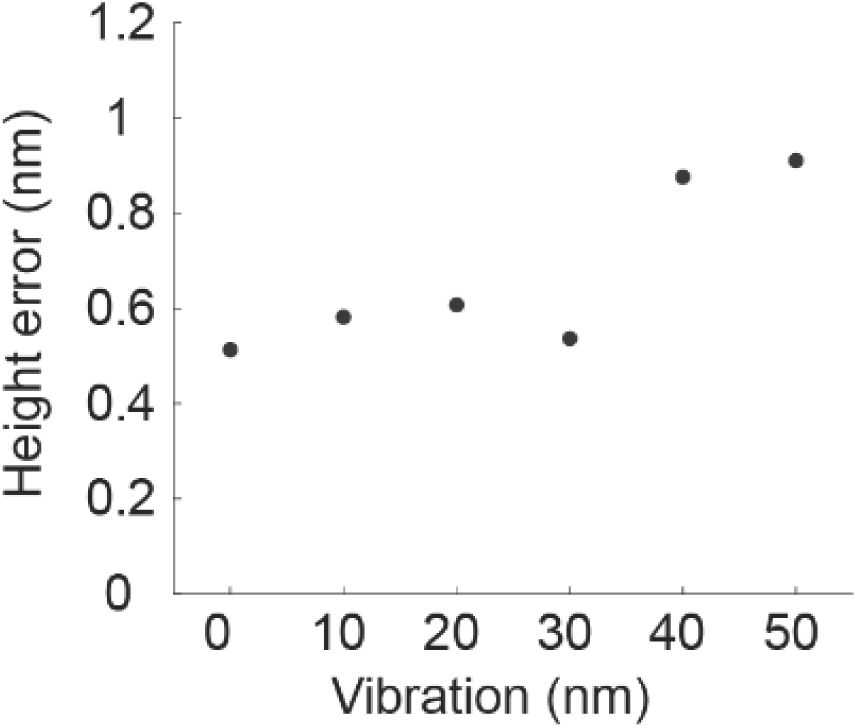
Effect of acoustic vibration on CSOP measurements. We simulated bead vibrations during image acquisition by overlaying several circles at different locations to construct an image frame. The height error was calculated as the standard deviation of heights measured in 25 independent simulations.

**Table S1:** Resolution limit of CSOP. Comparison of localization error between STORM and CSOP measurements using experimental measurements and simulations. A 6.8μm lipid coated bead containing 1,000/μm^2^ DOPE-Liss Rhod B was made and the labeling densities of 42/μm^2^ and 233/μm^2^ were emulated by reducing the imaging laser power. The predicted or measured error at labeling density of 550/μm^2^ are predicted based on the linear fitting of the simulated or experimental data. The camera signal was converted to number of photons based on its quantum yield, 0.82. Peak photon is the number of photons detected at the brightest pixel, and SNR is the ratio peak signal to photon shot noise. The predicted CSOP error is obtained by simulation (Materials and Methods). Although STORM can localize in 3D, the error indicated in the table represents 2D localization. CSOP localizes in 1D.

|  | STORM | CSOP |  |  |  |
| --- | --- | --- | --- | --- | --- |
| Labeling density | Single molecule | 42/$\mu\text{m}^2$ | 233/$\mu\text{m}^2$ | 550/$\mu\text{m}^2$ | 1,000/$\mu\text{m}^2$ |
| Peak photon | ~400 | 30 | 168 | 397 | 720 |
| SNR | 20 | 5.5 | 13 | 20 | 27 |
| Total # photon | 3,000 | 39,000 | 218,400 | 515,000 | 936,000 |
| Predicted error | 4 nm (Ref. 1) | 1.4 nm | 0.48 nm | 0.35 nm | 0.28 nm |
| Measured error | 8 nm (Ref. 2) | 2.26 nm | 1.01 nm | 0.65 nm | 0.48 nm |

**Table S2:**
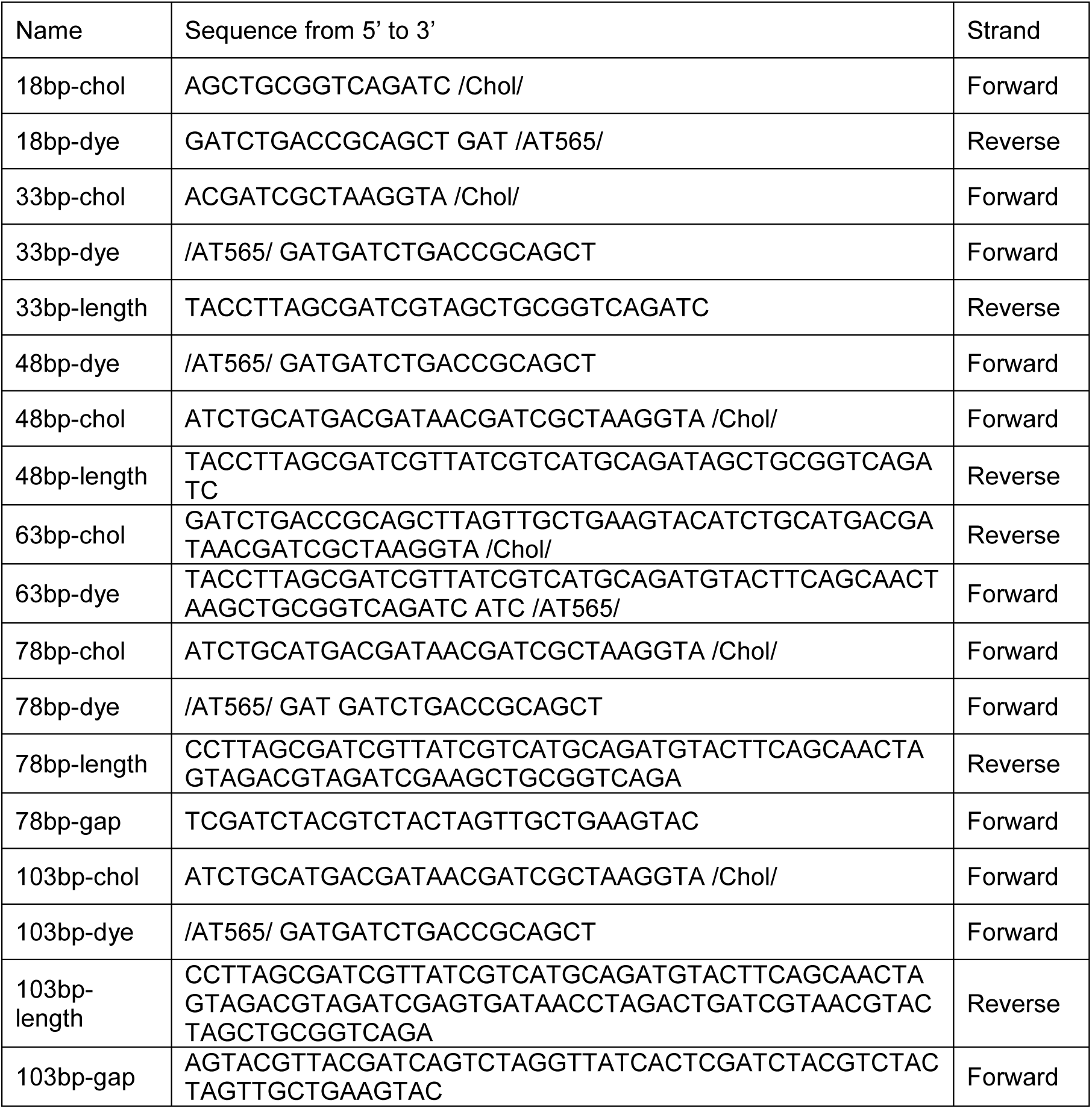
Design of size standard double-stranded DNA. Double-stranded DNA of varying size is assembled using one reverse strand and one or more forward strands that sequentially align onto the reverse strand without gaps. Forward strands contain an atto-555 dye at their 5-prime end and a cholesterol moiety at their 3-prime end, except for the 18bp version, which contains an atto-555 dye on the reverse strand.

**Movie S1. MD simulations of multi-domain proteins at large surface densities.**

MD simulation of 11-domain proteins with persistence length *L_p_* = 12 nm at surface density of ∼15000 chains / μm^2^.

**Movie S2. MD simulations of multi-domain proteins in crowding solution.**

MD simulation of 9-domain proteins with persistence length *L_p_* = 12 nm at dilute surface density of ∼200 chains / μm^2^. Spherical particles of diameter 5.25nm are added to the bulk of the simulation box at 5% w/v to model the effects of macromolecular crowding; these particles (shown in yellow) have been rendered slightly transparent for clarity.

**Movie S3. MD simulation of a protein-antibody complex.**

MD simulation of 8-domain proteins with persistence length *L_p_* = 12 nm at surface density of ∼10000 chains / μm^2^. The antibody is modeled as a rigid assembly of 5 spheres using typical dimensions of an IgG antibody. The red particle represents a Fab region on the antibody and binds to the terminal domain of the protein (shown as a yellow particle).

